# Programmable orthogonal and rapid sequential DNA strand displacement for fluidic-exchange-free highly multiplexed fluorescent imaging

**DOI:** 10.1101/2025.09.22.672701

**Authors:** Yanju Chen, Ryan N. Delgado, Ethan Xu, Constance L. Cepko, Fan Hong

## Abstract

Multiplexed fluorescent imaging methods are essential for studying cellular function by visualization of biomolecules in cells and tissues with high-resolution spatial information, but most suffer from limitations such as low multiplexity because of spectral overlap between the used fluorophores. Fluidic exchange based sequential multiplexed imaging overcomes this limitation but requires time-consuming incubation and washing steps. We introduce a novel multiplexed fluorescent imaging method that uses rapid, orthogonal DNA strand displacement reactions to enable unlimited multiplexed fluorescent imaging without fluidic exchange. The signal switching between targets is achieved by strand displacement with added non-fluorescent DNA displacer strands, no fluidic washing step is required for sequential multiplexed imaging. We experimentally screened a set of rapid and orthogonal DNA displacement sequences for probe design and applied it for RNA imaging, which takes less than 30 seconds per round to complete in fixed cells. Because of the vast sequence design space of DNA probes, theoretically unlimited multiplexity can be achieved. Using 25 developed rapid orthogonal probes, we achieved 25-plex RNA imaging in a single fluorophore channel in fixed cells within 20 minutes. To further demonstrate robustness and practical usage of DIRSE-based imaging, we showed 24-plex RNA imaging with the method in retinal tissues and resolved different cell types. This DIRSE mechanism significantly simplifies the high-plex fluorescent imaging process with pre-programmed DNA probes and has broad biotechnical applications for future medicine and diagnostics.

## INTRODUCTION

Fluorescence imaging of biological targets is critical to reveal the complex organization of biomolecules within and across cells to study cell functions and tissue organization^1-3^. Techniques such as fluorescence in situ hybridization (FISH) and immunofluorescence (IF) imaging, allowing direct visualization of targets in their intact cell and tissue environment, have been widely used to reveal critical details about the abundance and spatial arrangement of RNAs and proteins^4, 5^. To comprehensively study physiological states and biological functions of cells within their tissue context, high-dimensional molecular information must be resolved using high-plex fluorescence imaging to target numerous biomolecules. Consequently, advanced high-plex imaging techniques are highly desired to image multiple biological targets simultaneously within the same sample.

Multiplexed fluorescent imaging requires an unambiguous fluorescent signal switching between different targets. Conventional multiplexed fluorescence imaging uses different fluorophore-labeled probes to target multiple biomolecules, but the multiplexity is limited by spectral overlap of fluorophores, restricting multiplexing to 3–5 targets. Although hyperspectral properties of different dyes can be used for higher-plex imaging, more sensitive detectors, precise signal calibrations, and complex analysis algorithm are needed to resolve the signals from different dyes^6-8^. Recently, DNA-based barcoding and fluidic-exchange have enabled high-plex imaging by staining all targets with DNA-barcoded binding entities in one round, followed by iterative binding and washing of fluorescently labeled oligonucleotides (DNA imagers)^9-17^. By encoding signals combinatorially across multiple rounds of DNA imager exchange, the multiplexing was exponentially increased^18-24^. Fluidic exchange of DNA imagers has become the dominant signal switching mechanism for the current multiplexed fluorescent imaging with wide applications in neuroscience^25^, oncology^26^, and immunology^27-29^.

Despite its success, fluidic-exchange DNA imager methods face two major challenges: (1) buffer exchange steps can take tens of minutes to hours to have sufficient signal binding for the current target and thorough signal removal for last round targets, significantly slowing signal switching between rounds, and (2) the required fluidic devices are complex, costly, and prone to issues like air bubbles and tubing failures. Although commercially platform is available, they are generally very costly for acquisition and maintenance. These time-consuming washing steps, instrumentation complexities, and economic cost limit the widespread adoption of high-plex RNA imaging. To address these challenges, the recently developed DNA thermal-plex^30^ technique eliminated fluidic-exchange steps for multiplexed imaging by using temperature-controlled melting to sequentially and rapidly activate DNA thermal probes in situ. However, this method is limited to five thermal channels, profiling up to 15 targets when combined with 3 fluorophore channels. Additionally, it requires a specialized heating chamber and temperature control module, still adding instrumentation complexity. Thus, there remains a need for new signal switching mechanism to achieve simpler, faster, and more accessible high-plex fluorescent imaging.

We introduce a novel multiplexed imaging method that uses orthogonal and rapid sequential DNA <u>di</u>splacement reaction-based <u>s</u>ignal <u>e</u>xchange (DIRSE) mechanism to switch fluorescent signals between biological targets without fluidic exchange. DNA strand displacement allows a single stranded DNA invader to displace out a single stranded DNA from a DNA duplex through a toehold, with reaction kinetics being controlled through the DNA sequences^31^. In the DIRSE enabled multiplexed imaging, DNA probes are designed to encode fluorescent signals through the hybridization of an imager and a quencher strand, enabling them to bind targets in situ in a single step. The signal activation and removal are driven by non-fluorescent quencher and imager displacers added to the sample on-microscope, respectively. Displaced imager and quencher DNAs remain at low concentrations (picomolar), rendering their presence negligible in the imaging buffer and eliminating the need of washing with fluidic exchange. Displacement reactions are designed through simplified reaction pathways and optimal sequences to achieve rapid reaction rate for signal exchange between targets. Multiple rounds of signal switching can be achieved through consecutive additions of displacer strands, streamlining the workflow and reducing instrumentation complexity. Each round of signal switching is completed in under 30 seconds in fixed cells, and no special external accessories are required for on-microscope imaging beyond a standard fluorescence microscope and a general lab set-up. Because of the vast sequence space of DNA probes, experimentally unlimited multiplexity can be achieved for fluorescent imaging. We screened DNA displacement sequences from 144 different reactions and developed a set of 25 different DNA probes with rapid kinetics and high orthogonality, enabling 25-plex RNA imaging in fixed cultured cells within 20 minutes using a single fluorophore channel. We further demonstrated practical usage of the method in complex heterogenous tissues analysis by imaging 24 different RNAs in mouse retinal tissue to resolve different cell types.

## RESULTS

### Principle of multiplexed imaging with DIRSE

DIRSE leverages programmable, orthogonal, and rapid DNA strand displacement to activate and deactivate fluorescent signal of DNA probes in situ. In DIRSE method, the fluorescent signal of targets is encoded by pre-assembled DNA probes, which consist of a fluorophore-labeled imager strand and a quencher strand that hybridize to repress fluorescence. Signal activation and deactivation occur through rapid, DNA displacement reactions using non-fluorescent quencher and imager displacer strands, respectively. Multiplexed imaging is achieved through orthogonal and sequential displacement of reactions. Because of the vast sequence space of toehold DNA probes, DIRSE theoretically enables multiplexity of thousands of targets, without a limitation in multiplexity for experimental high-plex imaging^32^.

The core of DIRSE is the design of DNA probes that enable efficient fluorescence generation and removal (**Figure 1a**). Each DIRSE probe comprises an imager strand with three domains (b*, c*, d*) and a quencher strand with two domains (a, b). The two strands are hybridized via domain b/b*, allowing the quencher to repress the signal of fluorophore through direct quenching^33^. When applied for imaging of a biological targets, binding entities, such as in situ hybridization (ISH) probes and antibodies, with a barcode (domain c) bind to the specific targets, and the pre-assembled DNA probe hybridizes to the barcode via domain c*. This forms a complex of three strands on the targets: the barcode, quencher, and imager, with two single-stranded toehold domains (a on the quencher, d* on the imager) available for displacement. Fluorescent signal of imager is not observed because of efficient direct quenching^33^. All domains are designed to be 18 nucleotides long to ensure stable binding, vast sequence space, favorable thermodynamics, and rapid displacement kinetics.

**Figure 1.**
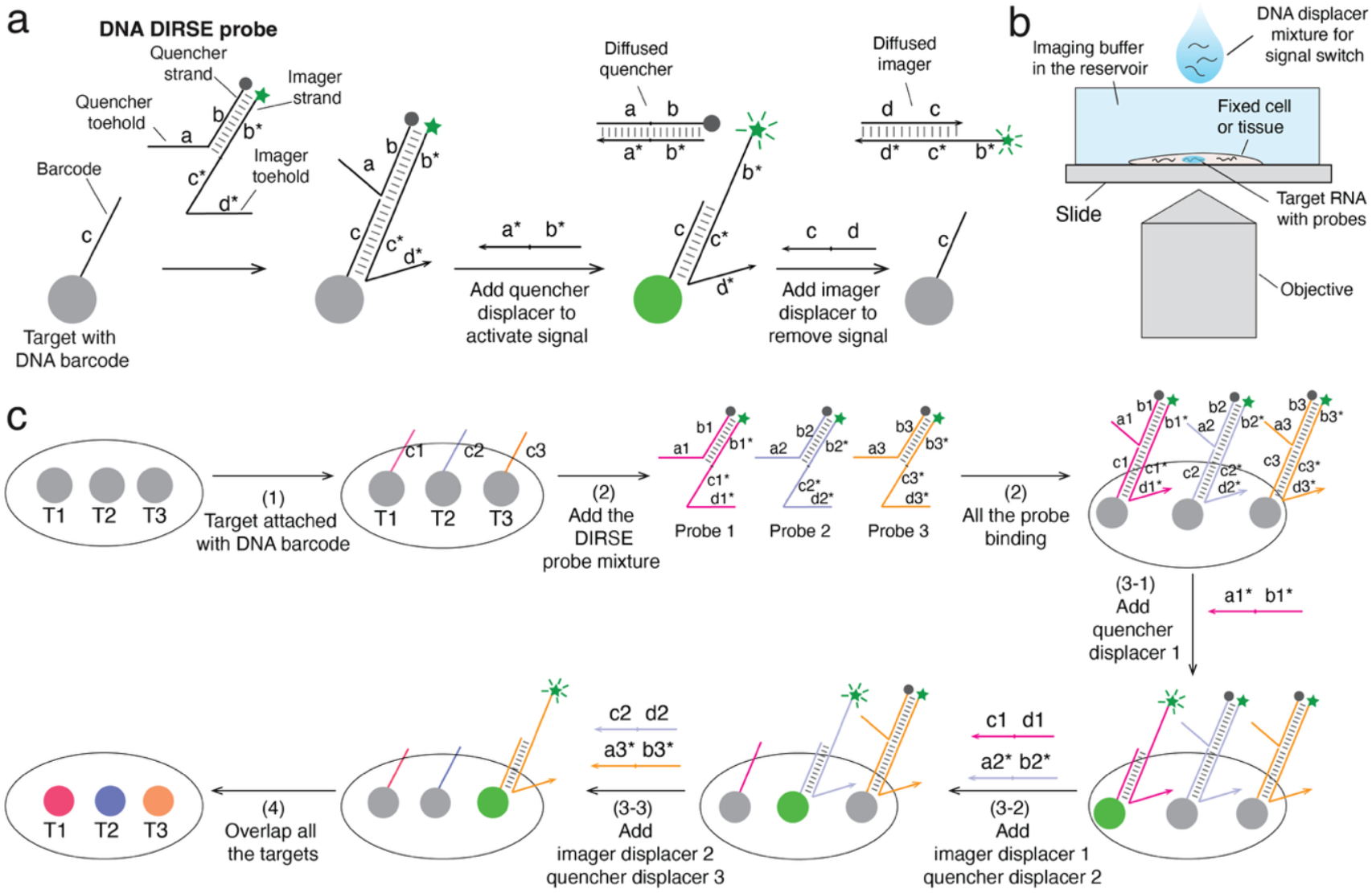
The scheme of DIRSE. (a) The designed molecular reactions for DIRSE. The RNA target hybridizes to the ISH probe, which carries a DNA barcode (domain c) that remains unhybridized. The DNA DIRSE probe is formed by the hybridization of an imager strand (domain ab) and a quencher strand (domain b^⋆^c^⋆^d^⋆^) through domain b/b^⋆^. The DNA probe binds to the target through the barcode (domain c), leaving single-stranded domains a and d^⋆^ as toeholds for signal activation and removal. To activate the signal, the quencher displacer initiates the strand displacement through the quencher toehold (domain a) and displaces the quencher from the target. Images can be acquired after signal activation. To eliminate the signal, the imager displacer will react with the imager to displace it from the target. (b) Scheme of the experimental setup. After the DNA probe binds to the target, the quencher and imager displacer mixtures are directly added to the sample with pipetting to initiate signal activation and removal. No external accessories were needed. (c) The scheme of multiplexed imaging with different rounds of DNA strand displacement reactions without washing steps exemplified with three targets. (1) Three different RNA targets (T1, T2, and T3) are labeled with three different barcodes (c1, c2, and c3); (2) Three different DNA DIRSE probes bind to their assigned RNA targets; (3-1) To activate the fluorescent signal for the first target, the quencher displacer 1 is added, followed by image acquisition; (3-2) The imager displacer for DNA probe 1 and quencher displacer for probe 2 will be added simultaneously to remove signal of T1 and activate signal of T2; (3-3) The imager displacer for DNA probe 2 and quencher displacer for probe 3 will be added simultaneously to remove signal of T2 and activate signal of T3; (4) After all the rounds of imaging for each target, the overall images for all the targets will be overlapped. All the displacer strands are non-fluorescent, and displaced imager and quencher were in extremely low concentration. No fluidic exchange is needed to wash away the access of displacers or displaced imagers and quenchers.

In the imaging process, non-fluorescent displacer strands are used to control fluorescent signal of targets in situ. As shown in **Figure 1a and 1b**, to activate fluorescence, a quencher displacer strand (domains b* and a*) initiates displacement via the quencher’s toehold domain a, releasing the quencher from the target into the imaging buffer and enabling the imager’s fluorescence. To deactivate fluorescence, an imager displacer strand (domains d and c) binds the imager’s toehold domain d*, displacing the imager off to the solution from the target in situ. As the displacer strands have no fluorescent label, they do not contribute to the background fluorescence when added to the sample. The displaced imager and quencher, present at picomolar concentrations, neither contribute to background fluorescence nor weaken the target fluorescent signal. Consequently, the fluidic exchange steps, which are typically required for probe change and background minimization, are eliminated.

For multiplexed imaging shown in **Figure 1c**, multiple orthogonal DNA probes bind distinct targets via barcodes on the biological binding entity. As displacer strands react only with their corresponding probes, parallel displacement reactions significantly reduce the number of displacement reaction rounds for multiplexed imaging. Except for the first and last rounds, imager displacers (for the previous round’s signal removal) and quencher displacers (for the current round’s signal activation) can be added simultaneously. The order of activation and deactivation is user-defined, and the resulting data can be reconstructed to produce high-plex RNA imaging profiles.

### Screening of DNA sequence with rapid displacement kinetics for DNA DIRSE probe designs

To achieve rapid signal switching between targets, DNA displacement reactions with fast kinetics are desired. Although the thermodynamic properties of DNA are well-characterized and modeled^34^, predicting the kinetics of DNA hybridization, particularly DNA strand displacement^35^, remains challenging. The kinetic rate of DNA strand displacement varies by orders of magnitude depending on their sequence^36^, and in situ strand displacement occurs in a fixed and complex cellular environment^37^, further complicating predictions. To identify DNA probes with rapid kinetics for signal switching, we experimentally screened 144 orthogonal DNA displacement reactions with different sequences (**Figure 2a, Supplemental Note 1, and Supplemental Figure S1**). ELAVL1 mRNA was used as the target to measure in situ strand displacement kinetics, and the corresponding ISH probes were designed using OligoMiner^38, 39^. To economically utilize a universal fluorescent imager for screening, a bridge strand was incorporated to assemble the universal imager onto the ISH probes (**Figure 2b**). Each bridge strand consists of two regions: one for binding the imager and another for validating the strand displacement reaction. Following the addition of displacer strand, the ELAVL1 mRNA signal decreased as the strand displacement reaction progressed. The time course fluorescent images were taken, and first-order kinetic fitting was performed to determine the rate constant for all the 144 reactions (**Supplemental Figure S1**). To ensure rapid kinetics to complete the reaction (95% of the reaction) within 30 seconds, only sequences with a kinetic rate greater than 1.2 × 10^5^ s^-1^ were selected for the design of DNA probes (**Figure 2c, 2d, 2e**). Ultimately, we identified 62 distinct sequences for rapid DNA strand displacement reactions. As each DNA probe’s signal activation and removal is achieved through two independent strand displacement reactions, 25 different DNA probes were designed with 50 DNA sequences from screened DNA displacement reaction with rapid kinetics for multiplexed imaging (**Supplemental Table S1)**. Because of the vast sequence space for the DNA probes, more kinetically rapid DNA probes can be further screened in necessary scenarios.

**Figure 2.**
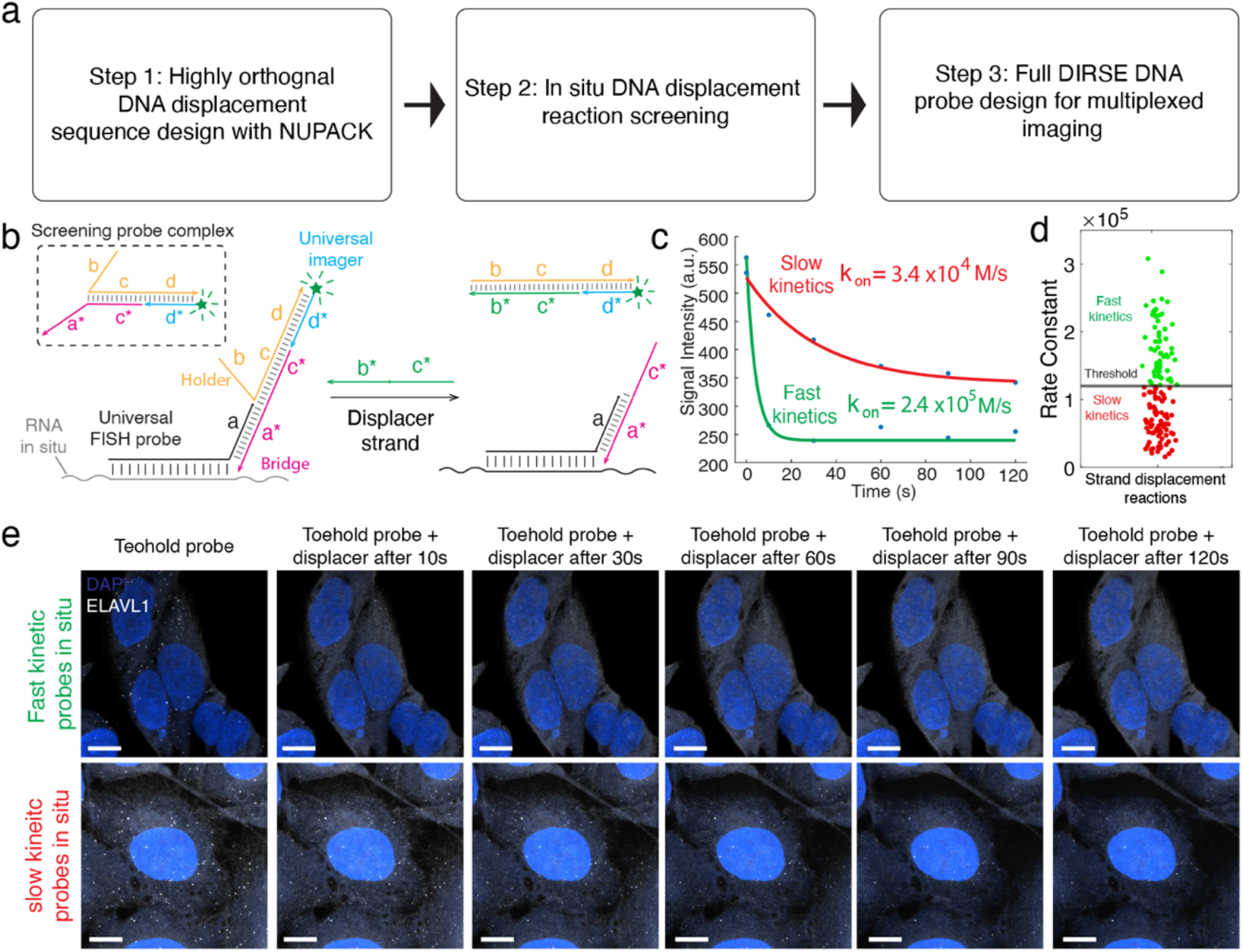
The screening of DNA displacement sequences with rapid kinetics. (a) The workflow for the rapid sequence screening to design DNA probes. (b) The reaction design to screen the DNA sequence for displacement reactions using a universal imager and ISH probe for Elavl1 mRNA. (c) The example of kinetic fitting for rapid (green) and slow (red) reactions. (d) The scattered plot for fitted rate constants of all 144 designed displacement reactions. Rapid and slow reactions are indicated in green and red colors, respectively. The threshold is 1.2 × 10^5^ s^−1^ to select the sequences for rapid displacement reactions. (e) The example fluorescent images for rapid and slow reactions at different time point. All the scale bars are 10 μm.

### Validation of rapid signal generation and removal of DIRSE in fixed cells

We next validated the two-step strand displacement reactions for signal activation and removal of the DNA probe by adding quencher and imager displacer DNAs to the sample by imaging RNA targets in cultured U2OS cells (**Figure 3a**). ELAVL1 mRNA was used as the target (**Supplemental Table S2)**, pre-assembled DNA probes were added to the sample to bind to the ELAVL1 ISH probe via the barcode. The imager is labeled with Atto-647 dye at 5’ end, and quencher is labeled with BHQ-3 quencher at 3’ end. After washing off the excess probes, the sample was placed on the microscope stage. Images were captured prior to the addition of any displacers to confirm that the DNA probe’s fluorescent signal was fully quenched (**Figure 3b**). To activate the DNA probe signal, a standard pipette was used to introduce the quencher displacer into the sample and make a final quencher displacer concentration to be 1 µM. Time-course recordings of signal change following the addition of quencher displacers were conducted to evaluate the speed of signal activation with a spinning disk confocal microscope. As shown in **Figure 3c, 3g and 3h**, the fluorescent RNA puncta showed up clearly after the displacement reaction, and the signal reached its maximum within 30 seconds. Similarly, the kinetic profiling was also performed after the addition of imager displacer after the quencher displacement reaction. The RNA fluorescent signal decreased rapidly within 30 seconds (**Figure 3d, 3i, and 3j**). Standard single molecule FISH (smFISH) experiment was conducted with a sample containing only the imager serving as the control to evaluate the quantities resolved by the DIRSE (**Figure 3e, 3f**). As shown in **Figure 3k**, the single-cell RNA expression of ELAVL1, measured using smFISH and DIRSE, is at the same level, confirming the accuracy of DIRSE for quantifying RNA gene expression. Negligible RNA was detected when the DNA probe is inactive or after the imager is displaced.

**Figure 3.**
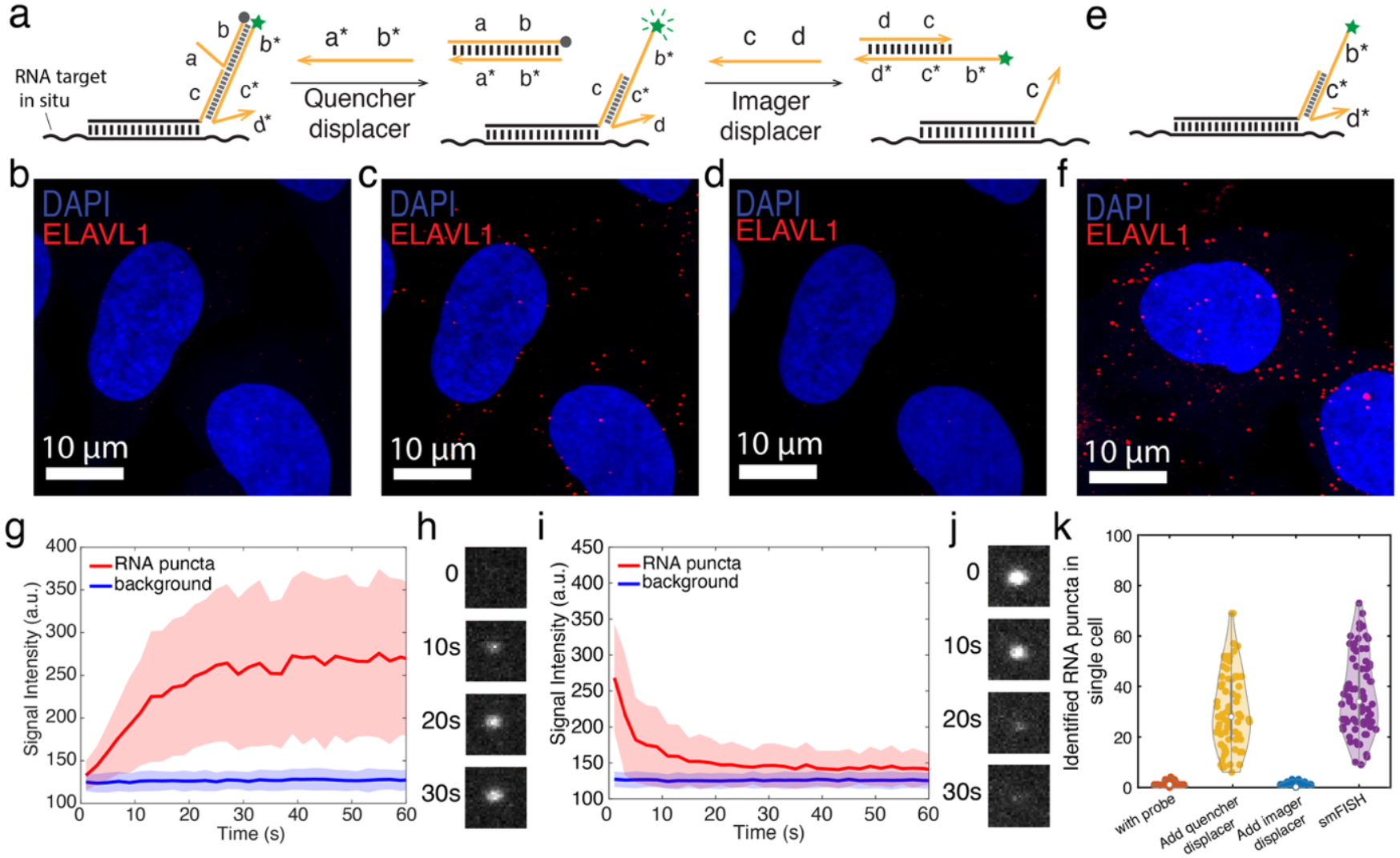
Validation of rapid signal activation and removal of DIRSE. (a) The scheme of RNA signal activation and removal of DIRSE. The added DNA displacer displaces out the quencher and imager DNA from the DNA probe bound with ISH probe on RNA targets to activate and remove signal, respectively. (b)(c)(d) The fluorescent imaging with DNA probe binding (b), after addition of quencher displacer (c), and after addition of imager displacer (d). (e)(f) the scheme of smFISH imaging and fluorescent images as positive control. (g)(i) The time course measurement of averaged RNA signal change after the addition of quencher displacer and imager displacer. Shaded regions denote mean ± s.d from all the RNA puncta in the field of view. (h)(j) Example images of a single RNA in situ at different time points after the addition of quencher displacer and imager displacer, respectively. The dimension of the images is 1 μm × 1 μm. (k) The violin plots of RNA puncta count after the DNA probe binding, addition of quencher displacer, addition of imager displacer, and smFISH positive control. The cell number used to count RNA copy number per cell for the samples were 58, 74, 61, and 73, respectively.

### Rapid 25-plex cellular RNA imaging using DIRSE in single fluorophore channel

We then proceeded to validate 25-plex RNA imaging using the designed 25 different DNA probes. 25 distinct RNAs, ranging from low to high expression levels, were selected based on bulk RNA sequencing of U2OS cells^40^. All imagers of DNA probes were labeled with Atto-565 dye, and all quenchers were modified with BHQ-2. Therefore, only the 565nm fluorophore channel was used for the 25-plex imaging to demonstrate the robustness of the DIRSE imaging. OligoMiner was used to design the ISH probes, with 25 unique barcodes appended to their 3’ ends for corresponding DNA probes (**Supplemental Table S2)**. The probe’s binding with barcode was analyzed in silico at 37 °C (**Supplemental Figure S2**). The 25 different DNA probes were first applied to image their assigned RNA targets individually to confirm their rapid kinetics (**Supplement Figure S3**). The fluorescent signal of all the DNA probes can be activated and removed within 30 seconds (**Supplemental Figure S3**). We further tested the orthogonality of 25 different DNA probes, only the corresponding displacer can activate or remove the fluorescent signal (**Supplemental Figure S4**).

To achieve 25-plex RNA simultaneously imaging, all the ISH probe hybridization were applied to the fixed cell for overnight hybridization. A mixture of the pre-assembled 25 DNA probes was then added to bind their respective RNA targets in fixed U2OS cells in a single step. To sequentially activate and removal the DNA probe signal through 25 rounds of displacement reaction, a set of 25 displacer mixtures, each at a concentration of 100 µM, was prepared. As shown in **Figure 4a**, the displacer for first round contained only the quencher displacer for DNA probe 1, the displacer for the final round contained only the imager displacer for DNA probe 25, and the displacer mixtures in the middle round N contained quencher displacer for DNA probe N and imager displacer for DNA probe N-1. The displacer mixtures were sequentially added to the sample to initiate strand displacement, activating the fluorescent signal for the current round and removing the signal from the last round. Each round of strand displacement lasted 30 seconds, followed by fluorescent image capture. The 25-plex RNA images were acquired using a single fluorophore channel at 565 nm in less than 20 minutes (10 seconds for operation, 30 seconds for displacement reaction, 5 seconds for imaging per round) for a single field of view.

**Figure 4.**
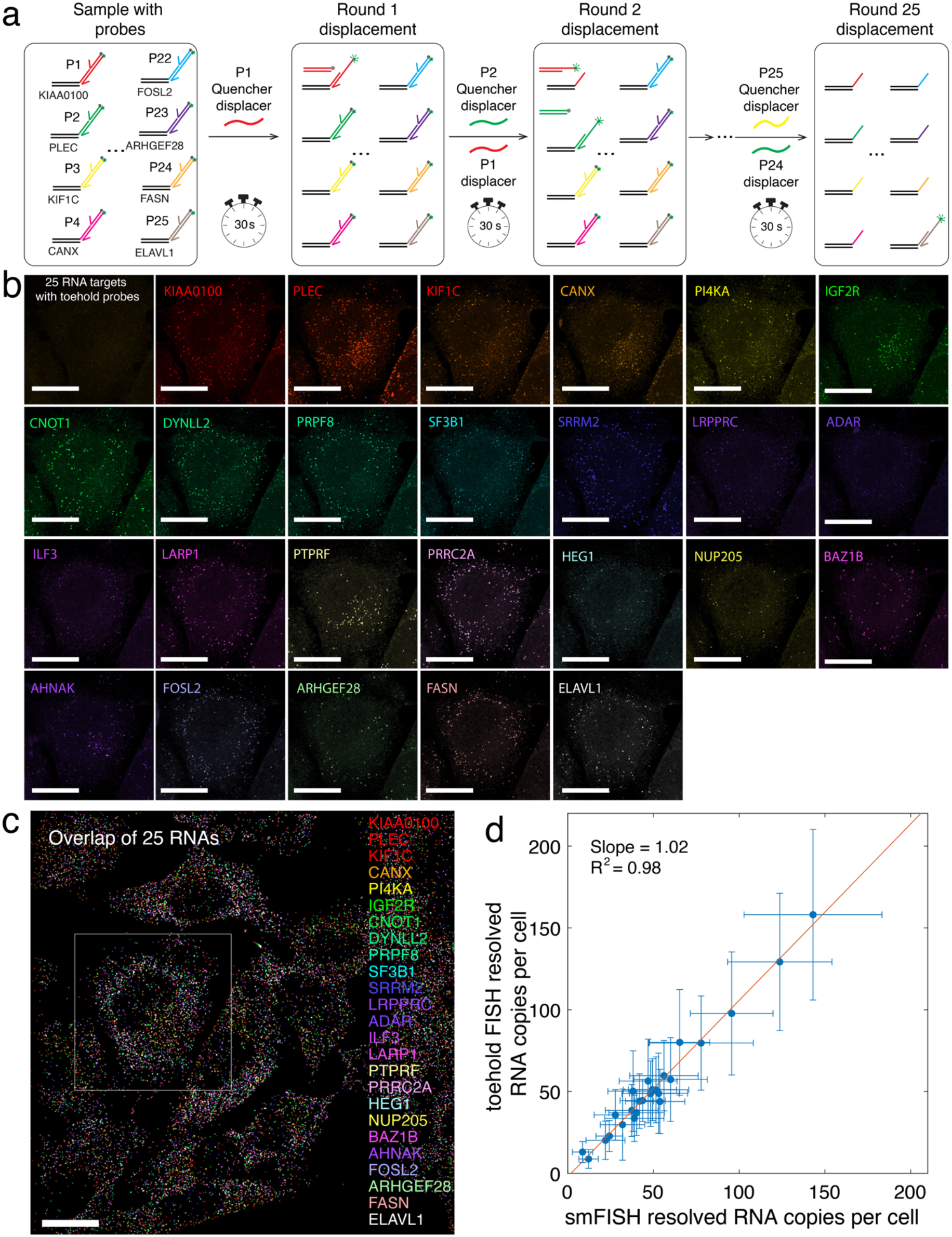
Rapid 25-plex RNA imaging with DIRSE in fixed cells in a single fluorophore channel. (a) Schematic of the multiplexed imaging process using sequential strand displacement reactions. All 25 DNA probes are pre-assembled and labeled with Atto-565 fluorophores. The 25 distinct ISH probes are hybridized with their RNA targets in situ, followed by the application of the25 DNA probes for binding in one step. Quencher and imager displacer are sequentially added to the sample to complete the strand displacement for their corresponding RNA’s signal activation and removal. Fluorescent images were captured after each round of strand displacement. (b) Individual fluorescent images for the 25 RNA targets after each round of strand displacement in the white boxed region in (c). All the scale bars are 20 μm. (c) Composite images showing the 25-plex RNA overlap. (d) Single-cell RNA expression comparison between DIRSE (n_DIRSE_ = 44 cells) and smFISH (n_smFISH_ > 30 cells for the statistics for all the 25 RNA targets) for 25 RNA targets. Error bars indicate the standard deviation of analyzed cells.

After the registration of all the images across 25 rounds based on nucleus staining with DAPI, the fluorescent images of 25 RNAs in a single representative cell were shown in **Figure 4b**, clearly depicted all 25 RNA species with each RNA name and assigned DNA probes. The overlap of 25 RNAs in a whole field were shown in **Figure 4c**, with a white boxed region indicating the cell in **Figure 4b**. The cells were segmented and all the 25 RNAs were identified to obtain single-cell gene expression profile from the fluorescent images. The smFISH fluorescent images were collected for each individual RNAs (**Supplemental Figure S5**) to obtain their single cell expression level to compare with the 25-plex DIRSE imaging. As shown in **Figure 4d**, RNA expression levels obtained from DIRSE correlated strongly with those from individually validated smFISH imaging with a slope of 1.02 and correlation of 0.98, confirming that high-plex DIRSE reliably quantifies RNA in cultured cell samples.

Compared to fluidic-exchange-based multiplexed RNA imaging, DIRSE significantly reduced signal switching time from tens of minutes to 30 seconds. The time consumption of overall multiplexed imaging workflow consists of three parts, (a) signal switching between targets, (b) sample imaging time, (c) microscope stage moving between different field of views. To quantitatively compare the time consumption of DIRSE and fluidic-exchange workflows, we calculated the overall time consumption as a sum of the three parts under different rounds of imaging and size of imaging area (see more details in **Supplemental Note 2**). Generally, the larger the number of exchange rounds is, the faster the DIRSE workflow is compared to the fluidic exchange workflow. The conditions under which the DIRSE workflow is 2-fold, 5-fold, 10-fold, 20-fold, and 30-fold faster are indicated as red lines in the heatmap (See **supplemental Note 2** and **Figure S6**).The cost of DIRSE is also low because no additional accessories are required beyond a standard fluorescence microscope and DNA probes (see more details in **Supplemental Note 3**).

### Validation of DIRSE in mouse retinal tissues

We further validated DIRSE in complex heterogenous tissue samples using PFA-fixed mouse retinal tissue. PRKCA mRNA, which serves as a cell-type-specific RNA marker for rod bipolar cells located in the inner nuclear layer of retinal tissue (**Figure 5a**), were used as the target. The ISH probe was designed with a DNA barcode appended at its 3’ end for DNA probe binding (**Supplemental Table S3)**. After fixation and permeabilization of the retinal tissue, the ISH probe was incubated overnight to bind PRKCA mRNA, followed by the incubation with the DNA probe that hybridizes to the barcode. The control sample was treated with only the imager binding for smFISH imaging. Before adding the quencher displacer to activate the signal, minimal signal was observed from the retinal tissue sample, indicating high quenching efficiency (**Figure 5b**). After adding the quencher displacer strand, time-course measurements of fluorescent signal were performed for all RNAs in the field of view. As shown in **Figure 5c**, most of the signal for PRKCA mRNA showed up in the desired inner nuclear layer of the retinal tissue. The averaged time course fluorescent signal for all the RNAs increased and reached a plateau in approximately 60 seconds (**Figure 5e**), indicating reaction completion. Following the addition of the imager displacer, the RNA signal gradually decreased and became undetectable after about 60 seconds, confirming complete signal removal (**Figure 5d and 5f**). Compared to fixed cultured cells, the displacement reaction in retinal tissue was slower, likely due to the extracellular matrix and high cell density. Retinal tissue is known for its high cell density^41^, which facilitates external signal reception and transmission to the brain. The rapid signal switching within 1 minute in retinal tissue demonstrated DIRSE’s broad applicability to tissue samples. The resolved single cell RNA copy number of PRKCA with DIRSE is at the same level as the one resolved by smFISH, indicating reliability of getting quantitative expression information in tissue samples (**Figure 5g**).

**Figure 5.**
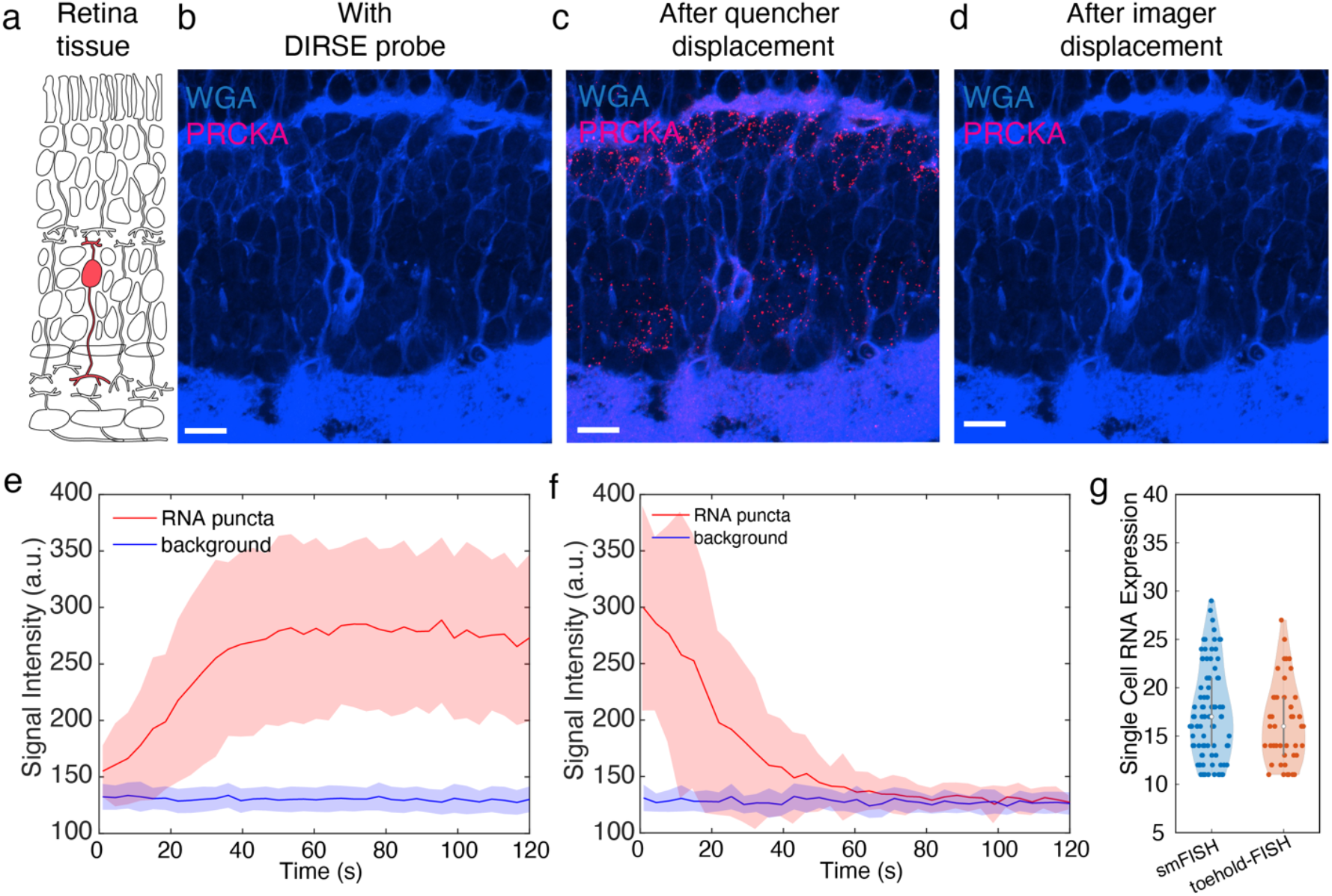
RNA imaging with DIRSE in mouse retina tissue. (a) The structures of retina tissue and selected Prcka mRNA markers located at the internuclear layer of the retinal tissue. (b)(c)(d) The fluorescent images for Prcka mRNA imaging after the DNA probe binding, quencher displacement, and imager displacement. The cell membrane was stained with WGA. All the scale bars are 10 μm. (e)(f) The time course measurement of single molecule RNA signal generation after the addition of quencher displacer and imager displacer. The reaction can be completed within 60s. Shaded regions denote mean ± s.d. with all the RNA puncta in the field of view. (g) The resolve single cell RNA copy number of Prcka mRNA with smFISH and DIRSE in retinal tissue samples. No significant difference is found between the two methods (n_sm-FISH_ = 79 cells, n=_DIRSE_ 45 cells).

### 24-plex imaging with DIRSE in mouse retinal tissues

The robustness of DIRSE in tissue samples supports its application for high-plex RNA imaging to resolve single cell types in complex tissue samples. The retina consists of ∼130 different cellular subtypes which are categorized into 7 different cell classes. This immense cellular diversity is arranged histologically into three distinct layers and work together to extract and process different visual features (**Figure 6a**). For instance, bipolar cells are a class of interneuron located in the inner nuclear layer and pass information from the photoreceptors in the outer nuclear layer to the retinal ganglion cells in the ganglion cell layer that transmit this processed information to the brain^42^. In mice, there are 15 different subtypes of bipolar cells which process different visual features and can be identified molecularly^43^. To demonstrate the DIRSE performance for cell typing in heterogenous and complex tissue structure, 24 different RNA markers for various cell types in the retina (see **Supplemental Table S3**) are selected for 24-plex DIRSE imaging. Oligominer was used to design ISH probe for all the RNAs and the barcodes for 24 different DNA probes are appended (see **Supplemental Table S3**). After the primary ISH probe and DNA probe binding, the retinal tissue sample was placed on the microscope stage. WGA labeled CF405s dye were used to stain the cell membrane. The prepared 24 displacer mixtures were sequentially added to the sample for DNA probe signal activation and removal, and imaging was taken after 1 minute of reaction for each round (**Figure 6b**). The whole workflow was completed within 30 minutes (10 seconds for operation, 60 seconds for reaction, 5 seconds for imaging per round) for the 24 rounds imaging to visualize 24 different RNAs for a single field of view.

**Figure 6.**
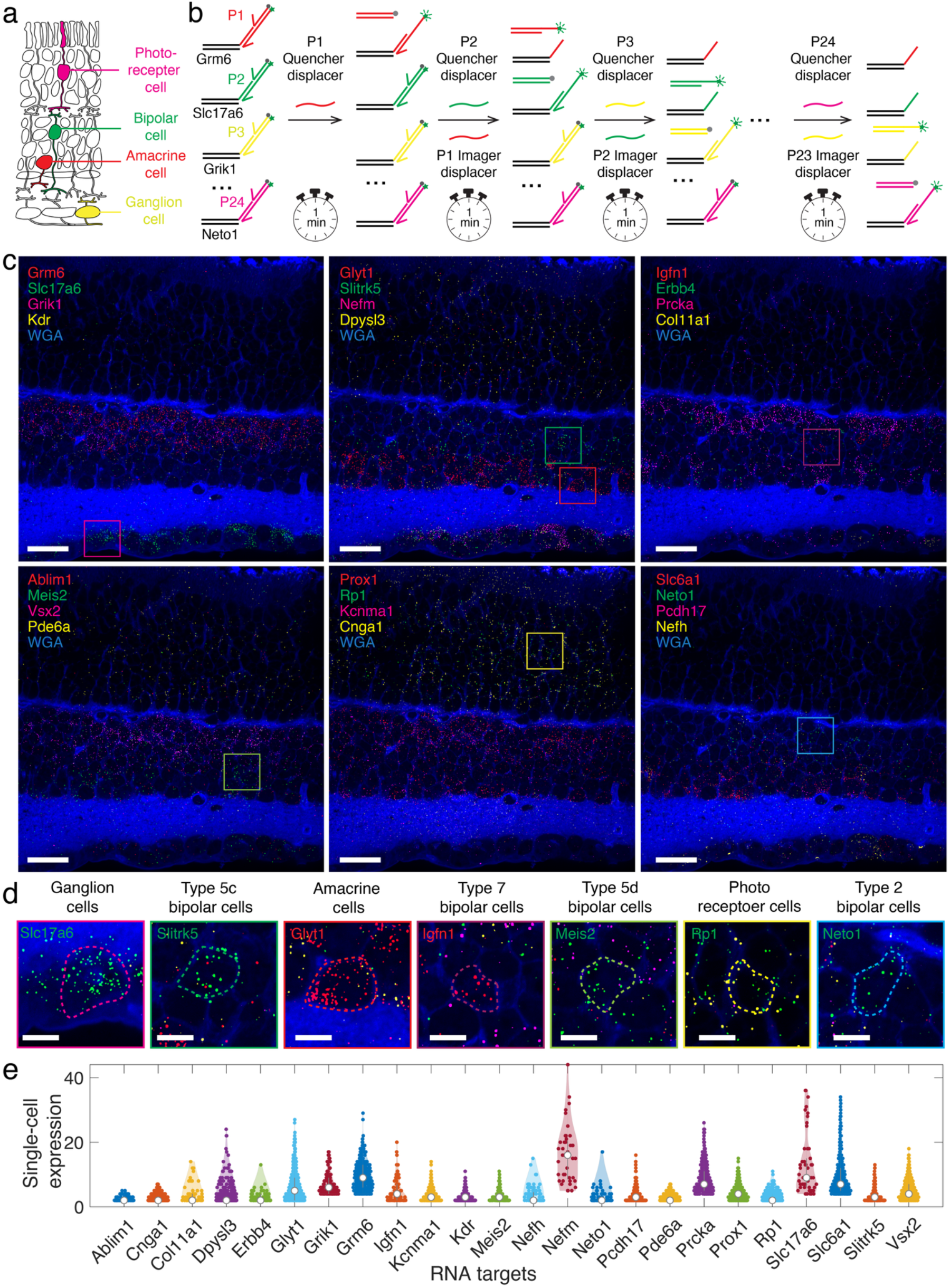
Spatial profiling of mouse retinal tissue with 24-plex RNA imaging in single fluorophore channel with DIRSE. (a) The organization of different cell types located in three different layers of retinal tissues. (b) The scheme of DIRSE for multiplexed imaging with 24 different DNA probes labeled with Atto-565 fluorophores; After all the DNA probe binding, non-fluorescent displacers are sequentially added to activate or removal RNA fluorescent signal via rapid orthogonal displacement reactions. (c) The fluorescent images of 24 RNAs in mouse retinal tissue after each round of DNA probe signal activation and removal for 1 min. The 24 RNAs were presented in 6 groups of overlapped images with 4 RNAs. The cell membrane was stained with WGA. The boxed regions in each overlapped panel are also shown in d. All the scale bars are 20 μm. (d) The selected resolved different cell types in the retinal tissues and corresponding RNA markers. All the scale bars are 5 μm. (e) The violin plots of single cell expression level of 24 RNAs. N > 40 cells were used for statistical analysis for all the RNA markers.

The individual 24 RNA images were shown in **Figure 6c** (see **Supplemental Figure S7** for individual fluorescent images of 24 RNAs). All the RNAs were clearly visualized in their desired region in the retinal tissue. Based on the 24-plex RNA expression, we were able to clearly resolve different cell types (**Figure 6d**), such as rod photoreceptor cells with Cnga1, amacrine cells with Glyt1, and ganglion cells with Nefm^44,45^. Several sub-type of bipolar cells in the internuclear region was also identified, such as type 2 bipolar cells with Neto2, type 5c with Slitk5, type 5d with Meis2, type 7 with Igfn1^43, 46^. We further compared the resolved single cell RNA expression data (**Figure 6e**) with smFISH (**Supplemental Figure S8**), confirmed consistency between the two methods for both spatial location and resolved single cell expression.

## DISCUSSION

The DIRSE mechanism enables highly multiplexed imaging in cells and tissues without fluidic exchange or additional accessories, utilizing rapid orthogonal strand displacement reactions. This approach significantly streamlines the multiplexed imaging workflow by reducing signal switching time from tens of minutes or even hours, typical in previous fluidic exchange-based methods, to less than one minute. It supports on-scope imaging without requiring costly or complex instrumentation. By simply adding displacer strands to the sample imaging chamber using standard pipettes, rapid signal activation and removal are achieved through pre-programmed rapid and orthogonal DNA displacement reactions.

As the number of signal exchange rounds increases, the fluidic exchange becomes more dominant in the overall workflow, and the multiplexed imaging process becomes significantly time-consuming and error prone. The workflow of DIRSE is increasingly faster compared with fluidic exchange when a higher number of exchange rounds is needed. We have developed 25 different DNA probes for 25 rounds of displacement reactions in a single fluorophore for 25-plex imaging. The vast DNA sequence space theoretically allows for the development of hundreds of such probes for unlimited multiplex imaging. Multiple spectrally separated fluorophores can be easily integrated with DIRSE to further increase the speed of the workflow.

The kinetics of DNA displacement reactions in situ not only depend on the sequence, but also depend on the target in situ biological environment. As the biological environment in cells and tissues is highly heterogeneous, DIRSE can be readily integrated with well-developed hydrogel embedding and tissue clearing^47, 48^ to further facilitate reaction speed for tissue imaging by creating a more homogenized reaction environment. Furthermore, these orthogonal and rapid DNA probes are not limited to multiplexed RNA imaging as we demonstrated here. They can be readily adapted for imaging other modalities by simply changing the binding entities that carry DNA probe barcodes, such as chromosomal DNA and proteins, in cell and tissue samples. DIRSE can be easily integrated with signal amplification, such as HCR, SABER, and RCA, to enhance its signal for biological imaging.

We have demonstrated DIRSE for multiplexed imaging in thin tissue sections, and it can also be applied to 3D deep tissues. Shorter DNA strands are shown to have fast penetration and hybridization kinetics within an hour in acrylamide hydrogel with millimeter dimensions^49^, significantly faster compared to days of fluidic exchange per round and potentially reducing the weeks of long signal exchange times in multiplexed thick tissue imaging^50, 51^. We envision that the DIRSE will profoundly change signal switching in multiplexed fluorescent imaging, significantly enhancing the accessibility and usability of multiplexed fluorescent imaging for broad applications.

## ACKNOWLEDGEMENT

This work is supported by the UF Start-up funding and UF Macknight Brain Institute Accelerator Award to F.H.

## AUTHORS CONTRIBUTIONS

Y.C. designed and performed the study, analyzed the data. R.N.D prepared the mouse retina tissue. E.X. performed the experimental study. C.L.C. contributed to the supervision of the study. F.H. conceived, designed, and supervised the study and wrote the manuscript. All authors edited and approved the manuscript.

## DATA AND MATERIALS AVAILABILITY

The main data supporting the results in this study are available within the main text and the Supplementary Information. The datasets generated during and/or analyzed during the current study are available from the corresponding authors on reasonable request.

