## supplemental information for "Programmable orthogonal and rapid sequential DNA strand displacement for fluidic-exchange-free highly multiplexed fluorescent imaging"

**Experimental Methods**

**Cell culture.** U2OS cells were maintained in DMEM medium (Gibco, catalog no. 10569) supplemented with 10%(v/v) fetal bovine serum (Gibco, catalog no. 10082), 100 U/mL penicillin, and 100 μg/ml streptomycin. The cells were cultured at 37 °C in the presence of 5% $CO_2$. For cell washing and passaging, 1× DPBS and 0.05% Trypsin (Gibco, catalog no. 25300) were used. U2OS cell lines (ACTT-HTB-96) were used in this study.

**DNA probe design.** All the sequence for DNA displacement reactions was generated using NUPACK. Custom MATLAB code was used to evaluate the orthogonality of the designed displacement sequences. Half toehold strand displacement reactions were designed and experimentally verified with rapid kinetics. After the kinetic screening, a sub-pool of sequence was identified with rapid kinetics. A pair of displacement reaction sequence from the sub-pool were assembled to design the DIRSE probes. The criteria to assemble the DIRSE probes is to minimize the imager secondary structures.

**RNA ISH probe design.** The gene accession IDs for selected RNA targets were collected and the sequence in FASTA file was downloaded from NCBI gene bank. Oligominer installed on a local computer was used to design the ISH probe with melting temperature setting between 41 ºC and 43 ºC and length setting between 32 and 38 nt. The generated probes were then aligned to hg38 or mus8 reference genome depends on the sample origin whether from human or mouse. After the ISH probe generation, DNA barcodes were appended to the 3' end to the DIRSE probe assigned.

**DNA and probe synthesis.** All desalted formed primary probes were purchased from Integrated DNA Technologies (IDT). All primary probes for each RNA transcript were combined in an equimolar mixture at 2 μM total in RNase- and DNase-free ultrapure water (Invitrogen, catalog no. 10977) and stored at −20 °C until use. The fluorophore-labeled imager strands and quencher strands were ordered with HPLC purification from Metabion. BHQ2 was used for Atto565 quenching, BHQ3 was used for Atto647 quenching.

**Cultured U2OS sample preparation.** 100 μL U2OS cells with a concentration of 1.5 $\times 10^5$ cells per milliliter were cultured on a chambered glass coverslip with 18 wells (Cat.No:81817) in an incubator (37 °C with 5% CO2) overnight. Culture media in the wells was aspirated, and the cells were rinsed twice in 100 μL 1× PBS. U2OS cells were fixed in 100 μL 1× PBS with 4% paraformaldehyde (v/v) (Electron Microscopy Sciences, catalog no. 191203) for 10 min and quenched in 100 μL 1× PBS with 100 mM $NH_4Cl$ for 5 min. After fixation, the cells were rinsed once in 100 μL 1× PBS and permeabilized in 100 μL 1× PBS with 0.5% Triton X-100 (Sigma, catalog no. SLBV4122) for 10 min. After permeabilization, the sample was washed twice in 100 μL 2× SSCT buffer (2× SSC buffer (Catalog number J60839.K2) with 0.1 % Tween-20 (Sigma, catalog no. SLCB0668)). 4× hybridization buffer was prepared by dissolving 40% dextran sulfate (Sigma, catalog no. 42867) in 8× SSCT buffer and stored at room temperature. For single-plex RNA target, ISH probe sets at 100 nM in total concentration were added to 100 μL hybridization solution containing 50% formamide (v/v) (Sigma, catalog no. S4117), 1× hybridization buffer and 10 U RNase inhibitor (Catalog number N8080119). For 25-plex assay, the ISH probe concentration was 500 nM. The sample was denatured at 60 °C for 3 min on a flat block thermocycler (Catalog number 4484078) and then incubate in a humid chamber at 42 °C overnight. After ISH probe binding, hybridization solution was aspirated. 100 μL 2× SSCT with 10% formamide (Sigma, catalog no. S4117) was prewarmed at 60 °C. Samples were then washed four times for 5 min each in 100 μL prewarmed 2× SSCT containing 10% formamide at 60 °C. Samples were rinsed once in 2× SSCT and then rinsed once in 1× PBS.

**DNA displacement sequence screening.** Screening solution was prepared by mixing 100 nM bridge strand, 120 nM holder strand, and 140 nM imager strand in 1× PBS (with excess holder and imager strand to ensure the assembly of three strands). 100 μL displacement sequence screening solution was added into the sample and incubate for 30 min. Samples were then washed twice for 5 min each in 1× PBS at 37 °C. Samples were then stained with 0.1 μg $ml^{-1}$ 4,6-diamidino-2-phenylindole (DAPI) in 1× PBS. After 5 min staining, the DAPI solution was aspirated, and the sample was rinsed in 1× PBS for 1 min. Finally, 100 μl of imaging buffer consisting of 1× protocatechuic acid (Sigma, catalog no. 03930590), 1× Trolox and 1× protocatechuic dioxygenase (Sigma, catalog no. 9029-47-4) in 1× PBS was added into the sample. 1 μL of displacer solution was added to the sample and mixed rapidly with a pipette, and fluorescent imaging was taken immediately for kinetic profiling.

**Retina tissue preparation.** The retina tissue fixation and section processed were described as previous assay[1]. The collected retina tissue with a thickness of 14 µM was embedded on a µ-slide 8 well high glass bottom (Cat.No:80807) and stored at -80 °C until use. 250 µL 1× PBS was added to wash the sample twice for 1 min. 250 µL 1× PBS with 0.5% Triton X-100 was used to penetrate the sample for 10 min. 250 µL 2× SSCT was then used to wash the sample for 1 min. 120 µL hybridization solution (50% formamide (v/v), 2× SSCT, 10% dextran sulfate, 12 U RNase inhibitor, primary probe) was added into the sample. For single-plex RNA target, the ISH probe concentration was 100 nM. For 24-plex assay, the total ISH probe concentration was 500 nM. The sample was denatured at 60 °C for 3 min on a flat block thermocycler and then incubated in a humid chamber at 42 °C overnight. The ISH probe washing step for retina tissues was performed as described for the cells, except that a volume of 250 µL solution was used. After DIRSE probe binding, 120 µL 10 µg $ml^{-1}$wheat germ agglutinin (WGA) conjugated to 405s (Biotium, catalog no. 29027) in 1× PBS was used to stain the cell membrane of the retina tissue. After 10 min staining, the WGA solution was aspirated. The sample was rinsed in 250 µL 1× PBS once and then adding 200 µL imaging buffer for imaging.

**Signal switching with DIRSE.** DNA displacer mixtures were prepared in a concentration of 100 µM in imaging buffer. When a round of imaging was completed and signal exchange was required for the next round, 1 µL of the DNA displacer mixtures was added to the sample and mixed by several repeated pipetting. After 30s (cell imaging) or 60s (retina imaging) incubation, the images were required again, and next round signal switching is performed repeatedly until all the round of displacement reaction completed.

**Microscopy imaging.** Images were acquired using a Nikon Eclipse Ti-E inverted microscope body equipped with a Yokogawa CSU-W1 spinning disk confocal unit, a Hamamatsu ORCA-FusionBT back-thinned camera, and a Plan Apo 100×/1.45 NA oil-immersion objective. Excitation was provided by 405 nm, 488 nm, 561 nm, and 647 nm solid-state lasers coupled into the spinning disk unit. Typically, 300ms exposure time was used for the imaging of both cultured cells and retinal tissues.

**Image analysis.** ImageJ was first used for the initial image processing, such as max z-stacking and channel split. Custom MATLAB (version 2023b) code was used for the image registration, RNA identification, and cell segmentation. In the scenarios of imager registration needed, the nucleus images in DAPI channel were used for image registration with MATLAB build in function. For RNA puncta identification, the fluorescent images were first filtered, and a signal threshold was applied to transform the images into a binary file. Another size threshold (12 pixel) was applied to identify the RNA puncta. The center of the RNA puncta was used as the location of the RNA. For cell segmentation, the cell body was manually drawn in MATLAB analysis pipeline and cell mask will be created for each cell segmented for the cultured U2OS cell samples. The segmentation of retinal tissue is performed with Cellpose[1] software, and the segmented cell boundaries were extracted for the downstream single cell analysis. For the cells not identified by the algorithm, manual segmentation was performed.

**Supplemental Note 1. Screening of DNA displacement sequence with rapid kinetics and high orthogonality.**

**1.1 Reaction design for kinetic screening.**

DIRSE imaging relies on rapid and orthogonal DNA displacement reactions, and each DIRSE probe undergoes two-step DNA strand displacement reactions to activate and remove the fluorescent signal, respectively. As it is challenging to precisely predict the kinetics of DNA displacement reactions in situ, we designed a pool of 144 different DNA displacement sequences and experimentally screened them for the final DIRSE probe designs, as shown in Figure 2a.

Since fluorophore-labeled DNA strands and in situ ISH probes are the most expensive reagents, synthesizing 144 fluorophore-labeled strands and ISH probes with different barcodes for experimental evaluation would be very costly. As shown in Figure 2b, we utilized a universal fluorescent imager (domain d*) and an ISH probe with a barcode (domain a) appended for the Elavl1 mRNA. A bridge strand (domains c* and a*), a holder strand (domains b, c, and d), and a displacer strand (domains c* and b*) were used to accommodate diverse displacement DNA sequences for economical sequence screening.

**1.2 Sequence design and fluorescent imaging data collection.**

NUPACK was used to generate a set of 144 different sequences for screening rapid DNA displacement reactions. All displacement reactions were tested with less than 10% crosstalk. All DNA probes and sequences were ordered from IDT. To evaluate displacement kinetics experimentally, a fluorescent DNA probe complex was assembled from the bridge strand, holder strand, and universal imager. Cultured U2OS cells were fixed and permeabilized for primary Elavl1 ISH probe hybridization, after which the assembled probe complex was applied to the fixed U2OS cells. After washing away excess probe complex, the corresponding displacer strand was added. Fluorescent images of RNA targets were captured using a microscope at 0 s, 10 s, 30 s, 60 s, 90 s, and 120 s after the addition of the displacer DNA strand.

**1.3 Data processing and kinetic fitting.**

RNA puncta were identified in the fluorescent images at 0 s, and their fluorescence intensity and location were recorded. The locations of the RNA puncta at 0 s were used to track their fluorescence signals in subsequent images at 10 s, 30 s, 60 s, 90 s, and 120 s at the same locations. The fluorescence signals of all RNA puncta were averaged at each time point, and first-order kinetic fitting was performed.

When the displacer DNA was introduced to the sample, the chemical reaction between the displacer DNA and the probe occurred as follows:

$$Displacer \; + \; RNA \cdot Imager \longrightarrow Displacer \cdot Imager \; + \; RNA$$

The differential equation for the reaction is:

$$\frac{d\,[RNA \cdot Imager]}{dt} \;=\; k_{disp}\,[Displacer]\,[RNA \cdot Imager]$$

Where $k_{disp}$ is the rate constant, the fluorescence signal observed through a fluorescence microscope is proportional to the [RNA-imager] concentration.

For a typical mRNA, the abundance is assumed to be 1–10,000 copies per cell, with the sample typically containing $10^4$ cells in a 100 µL imaging buffer chamber. Each mRNA is targeted by 48 unique probes. The concentration of bound imager is calculated to be $10^{-14}$ to $10^{-12}$ M. As the in-situ RNA-imager concentration (below pM levels) is extremely low compared to the displacer concentration (approximately 1 µM), the displacer concentration is assumed to remain constant throughout the reaction.

The integral of the differential equation above from 0s to time point t gave the following:

$$[RNA \cdot Imager] \;=\; [RNA \cdot Imager]_0\, e^{-k_{disp}[Disp]_0 t}$$

The initial $[RNA\text{-}Imager]_0$ concentration is proportional to the RNA puncta fluorescence signal at 0 s. The above equation was used to fit the data collected from fluorescence images at different time points, and the rate constant $k_{disp}$ was derived from the fitted curves. To ensure signal switching was completed within 30 seconds, with 96% of the signal activated or removed, a threshold for the rate constant was set for displacement sequence selection $1.2 \times 10^5\ s^{-1}$. Among 144 unique sequence designs, 62 reactions exhibited a rate constant above this threshold. Example kinetic fittings for DNA sequences with rapid and slow kinetics are shown in Figure 2c, and a scatter plot summarizing the rate constants from all 144 reactions is presented in Figure 2d. Corresponding fluorescence images at different time points are shown in Figure 2e.

**Supplemental Note 2. Comparison of total time consumption for the multiplexed imaging workflow between DIRSE and buffer-exchange based signal switching.**

In this section, we compare the overall time consumption of the multiplexed imaging workflow using DIRSE with fluidic exchange of DNA imagers. After sample preparation, the sample is typically placed on the microscope for multiplexed imaging. The multiplexed imaging workflow consists of three components: (1) signal exchange between different targets, (2) movement of the microscope stage across the sample, and (3) laser exposure and image acquisition.

1. **Signal switching time.**

The signal switching time for DIRSE per round, $t_{DIRSE}$, is 30 seconds, while the signal switching time for fluidic exchange, $t_{FE}$, is assumed to be 20 minutes[2, 3]. If n rounds of exchange are required, the total time for signal switching can be calculated as follows:

$$t_{switch-DIRSE} = (n-1)\, t_{DIRSE}$$

$$t_{switch-FE} = \ (n-1) t_{FE}$$

2. **The time used for the moving of microscope stage across the sample**

To image a large sample area, tiled imaging is required. Here, we assume a single slice of the sample is imaged along the z-axis. The microscope stage must move between tiles to cover the entire sample area. The sample has a dimension of D, with an imaging area of $D^2$. We assume the microscope is equipped with a camera providing 2304 × 2304 pixels at 6.5 µm per pixel and a 60× objective, typical setup for fluorescent RNA imaging. Thus, the dimension of a single field of view is approximately d = 249.6 µm. The total number of tiles required to cover the sample is calculated as follows:

$$N_{tile} = D^2/d^2$$

The average time for the microscope to move is $t_{microscope\text{-}step}$ = 1 second. The total steps that the microscope stage needs to move across all the samples is N-1. Consequently, the total time consumed for the microscope stage to move across the sample area is calculated to be:

$$t_{stage} = (D^2/d^2 - 1) \times t_{microscope-step}$$

3. **Camera imaging time**

When the microscope stage moves to each tile, images are taken by the camera for each tile. The camera imaging time depends on the channel number and exposure time. If three channels are used and exposure time for each channel is assumed to be 500 ms, and the camera imaging time at each tile is $t_{camera\text{-}step}$ = 1.5 s. The total camera imaging time is calculated to be:

$$t_{camera} = (D^2/d^2 - 1) \times t_{camera-step}$$

The difference between the DIRSE and fluidic exchange workflow is the signal switching time. Therefore, the overall total imaging time for a given sample with a dimension D for the two methods is calculated as the following:

$$t_{sample-DIRSE} = t_{switch-DIRSE} + t_{stage} + t_{camera}$$

$$t_{sample-FE} = t_{switch-FE} + t_{stage} + t_{camera}$$

The time consumption ratio of DIRSE to fluidic exchange workflow is calculated as:

$$Ratio = \frac{t_{sample-DIRSE}}{t_{sample-FE}}$$

The total time consumption depends on the sample size and the number of exchange rounds. We compared the time consumption ratio for sample sizes ranging from 0.1 to 20 mm and exchange rounds from 1 to 100. A heatmap of the time consumption ratio is presented in Supplemental Figure S6a. The conditions under which the DIRSE workflow is 2-fold, 5-fold, 10-fold, 20-fold, and 30-fold faster than fluidic exchange are shown in Supplemental Figures S6b, S6c, S6d, S6e, and S6f, respectively.

**Supplementary Note 3. Cost of DIRSE based RNA imaging.**

DIRSE based multiplexed imaging requires no additional instrument accessories, with all expenses tied to DNA probes. The cost of DIRSE based RNA imaging includes four components: a primary probe pool for RNA binding, DIRSE probe imagers, DIRSE probe quenchers, and displacer DNA strands.

A standard pool of ~48 unpurified probe oligonucleotides costs ~$200, yielding enough for 250 mL of 1 μM probe solution. An imager typically costs $200 for ~10 nmole, producing 1 mL of 1 μM solution. A quencher costs $80 for ~15 nmole, providing 1.5 mL of 1 μM solution. A displacer strand costs $5 for 10 nmole, resulting in 200 μL of 100 μM solution. In a typical DIRSE based RNA imaging experiment, ~120 μL of ISH probe at 100 nM is used, costing ~$0.012, while 120 μL of 200 nM DNA thermal probe costs ~$8 per experiment.

Costs can be lowered by ordering unlabeled DNA oligonucleotides and performing fluorophore and quencher conjugation in-house.

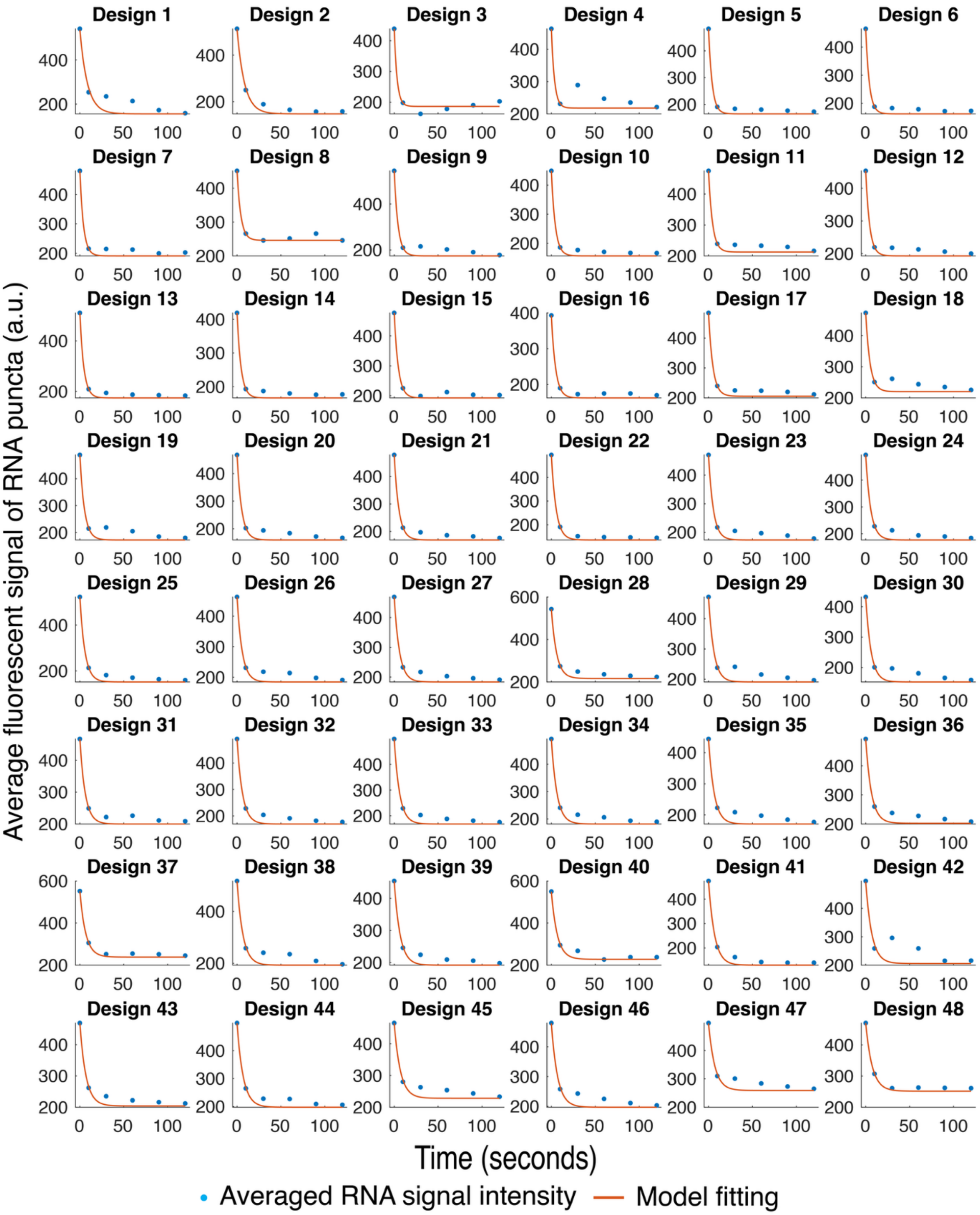

Design 1
Design 2
Design 3
Design 4
Design 5
Design 6
Design 7
Design 8
Design 9
Design 10
Design 11
Design 12
Design 13
Design 14
Design 15
Design 16
Design 17
Design 18
Design 19
Design 20
Design 21
Design 22
Design 23
Design 24
Design 25
Design 26
Design 27
Design 28
Design 29
Design 30
Design 31
Design 32
Design 33
Design 34
Design 35
Design 36
Design 37
Design 38
Design 39
Design 40
Design 41
Design 42
Design 43
Design 44
Design 45
Design 46
Design 47
Design 48
Average fluorescent signal of RNA puncta (a.u.)
Time (seconds)
Averaged RNA signal intensity
Model fitting

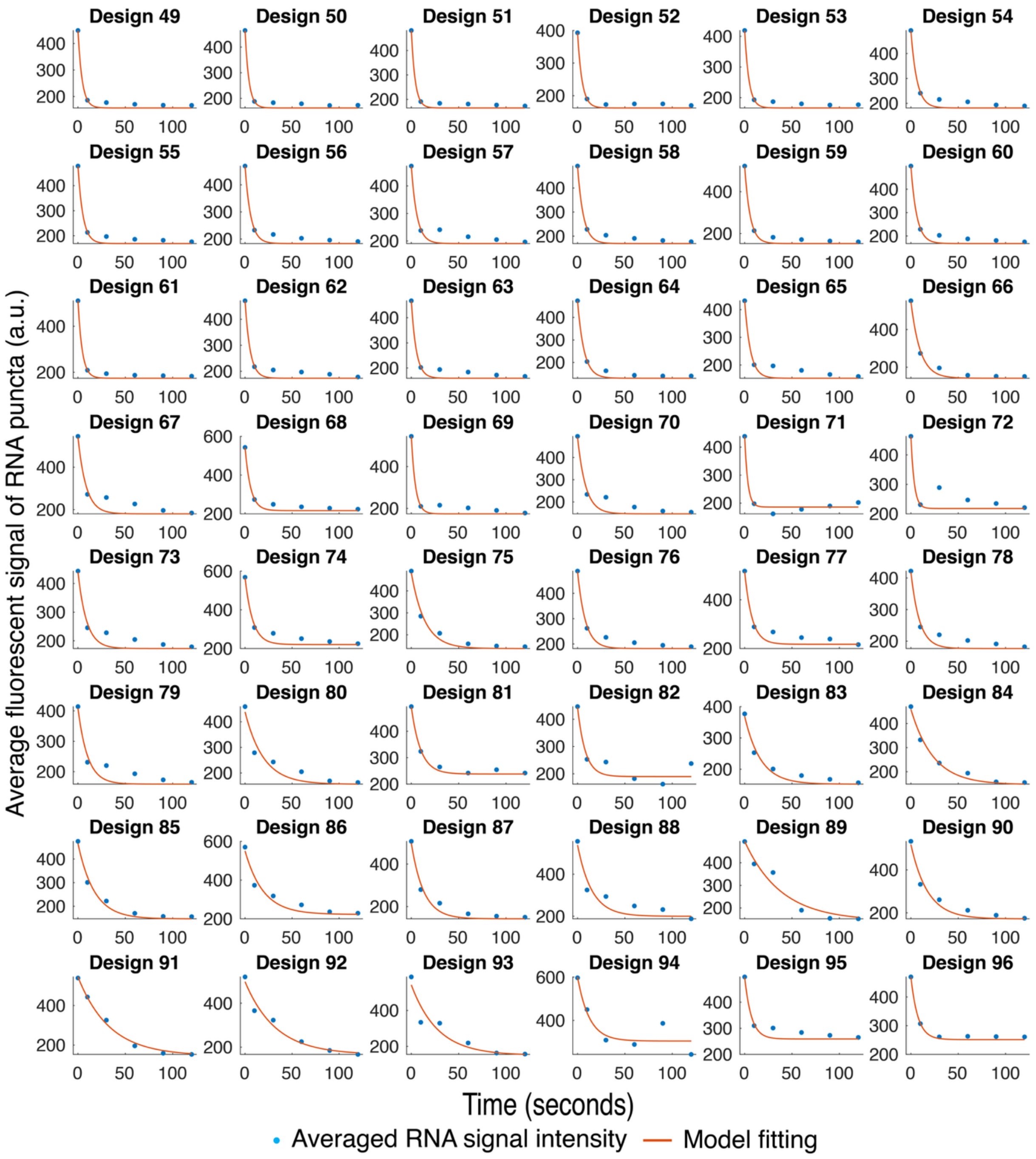
Design 49
Design 50
Design 51
Design 52
Design 53
Design 54
Design 55
Design 56
Design 57
Design 58
Design 59
Design 60
Design 61
Design 62
Design 63
Design 64
Design 65
Design 66
Design 67
Design 68
Design 69
Design 70
Design 71
Design 72
Design 73
Design 74
Design 75
Design 76
Design 77
Design 78
Design 79
Design 80
Design 81
Design 82
Design 83
Design 84
Design 85
Design 86
Design 87
Design 88
Design 89
Design 90
Design 91
Design 92
Design 93
Design 94
Design 95
Design 96
Average fluorescent signal of RNA puncta (a.u.)
Time (seconds)
Averaged RNA signal intensity
Model fitting

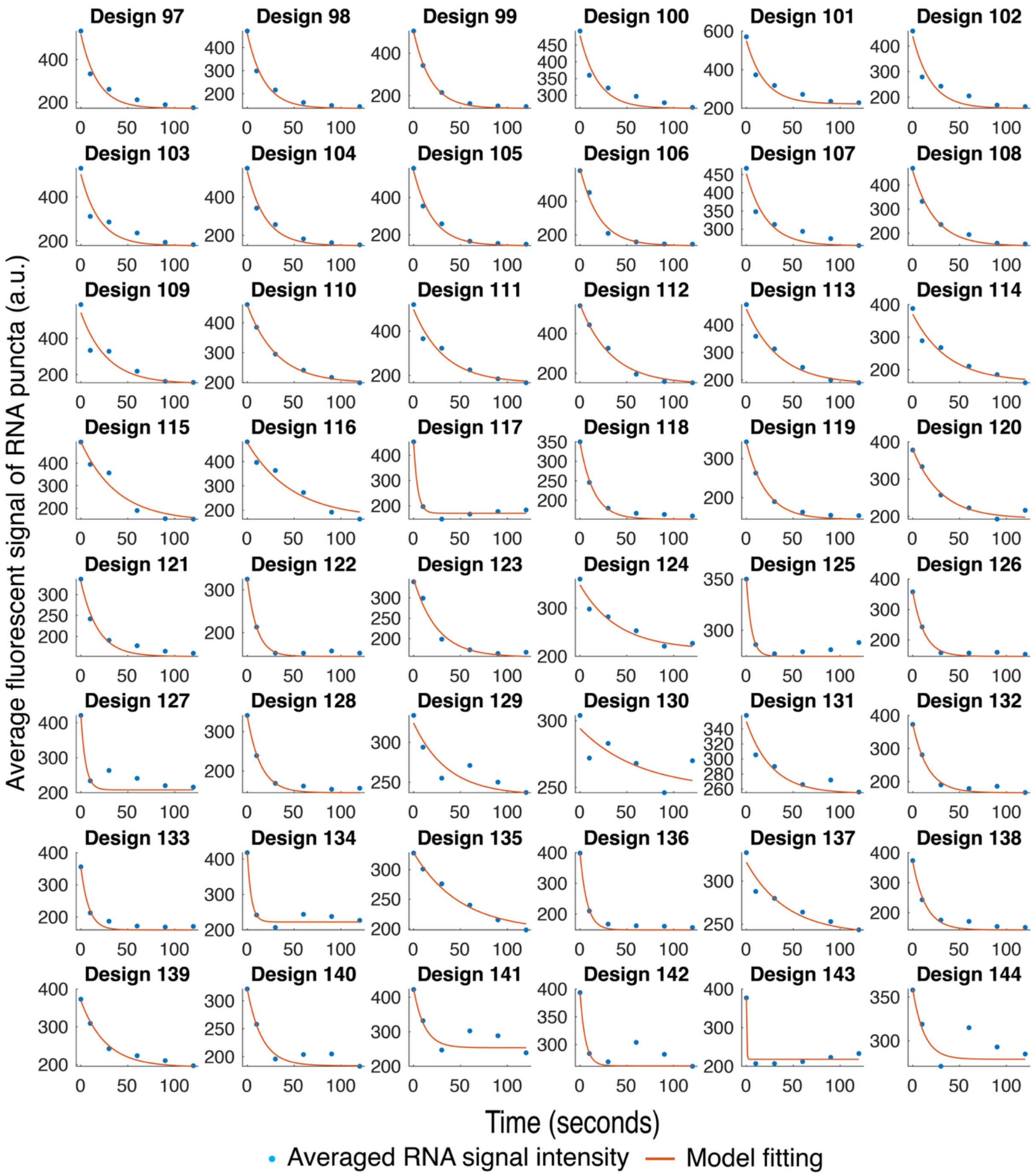


**Figure S1. The averaged RNA signal at different time point and kinetic fitting for all the designed 144 DNA displacement reactions for screening.**

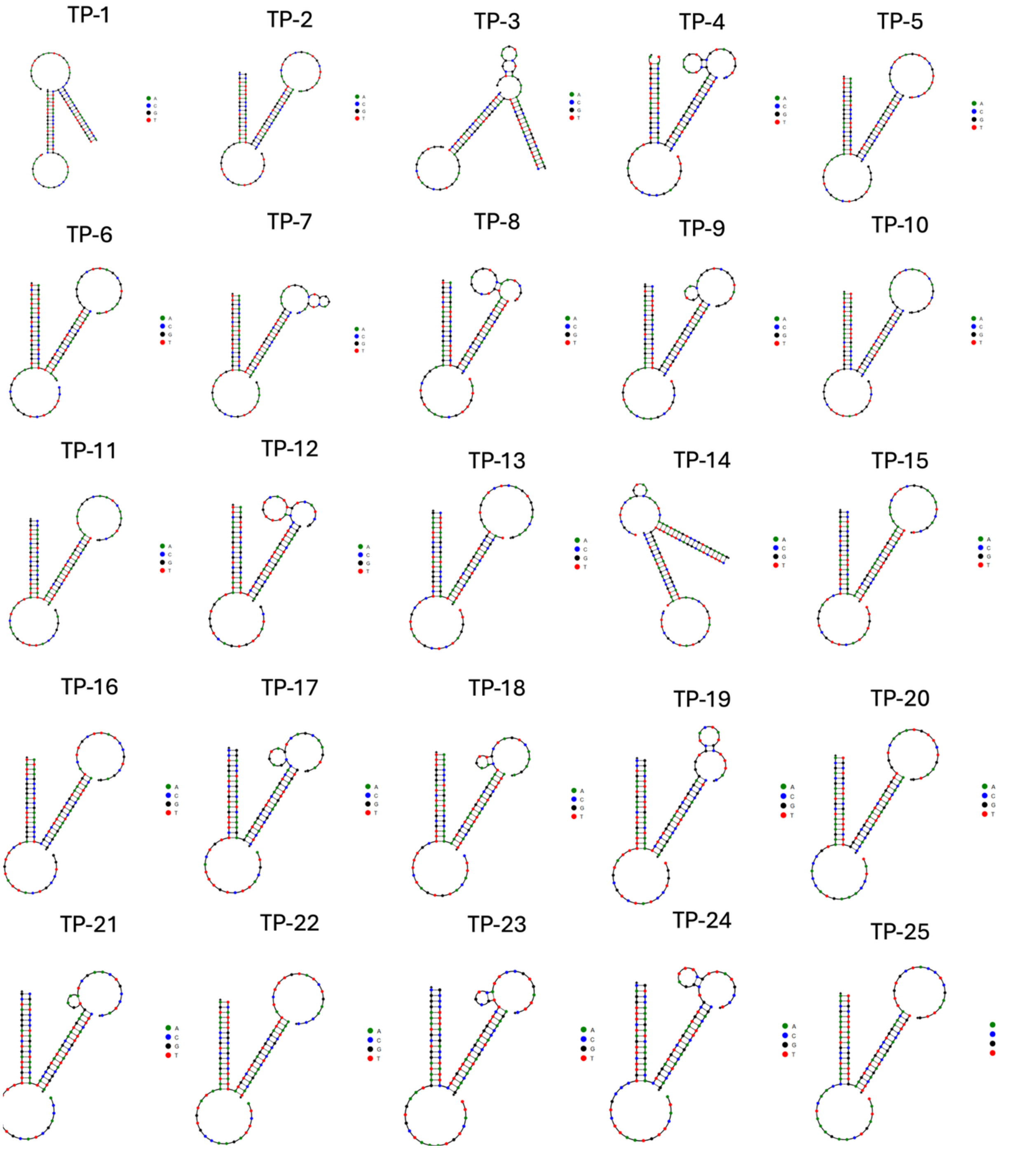


**Figure S2. The secondary structure analysis of probe-barcode complex.** NUPACK was used to analysis the minimal free energy state of the three-strand complex.

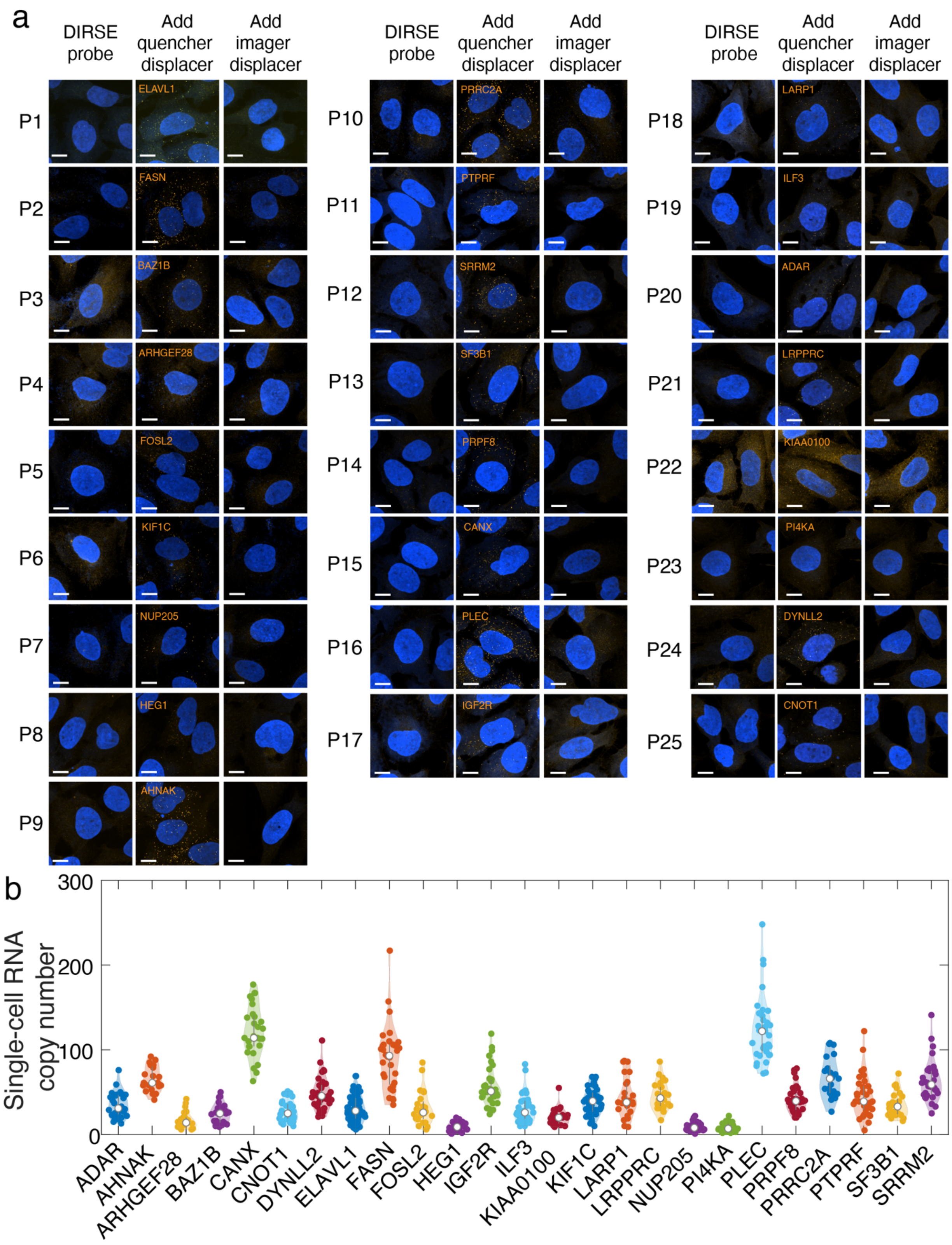


**Figure S3. The validation of 25 DIRSE probes in fixed cells.** (a) The fluorescent imaging of RNAs with DIRSE probes, after addition of quencher displacer, and after the imager displacers. Scale bars, 10 μm. (b)

The violin plots of resolved single cell RNA copy number after the DIRSE probe signal activation with the quencher displacer.

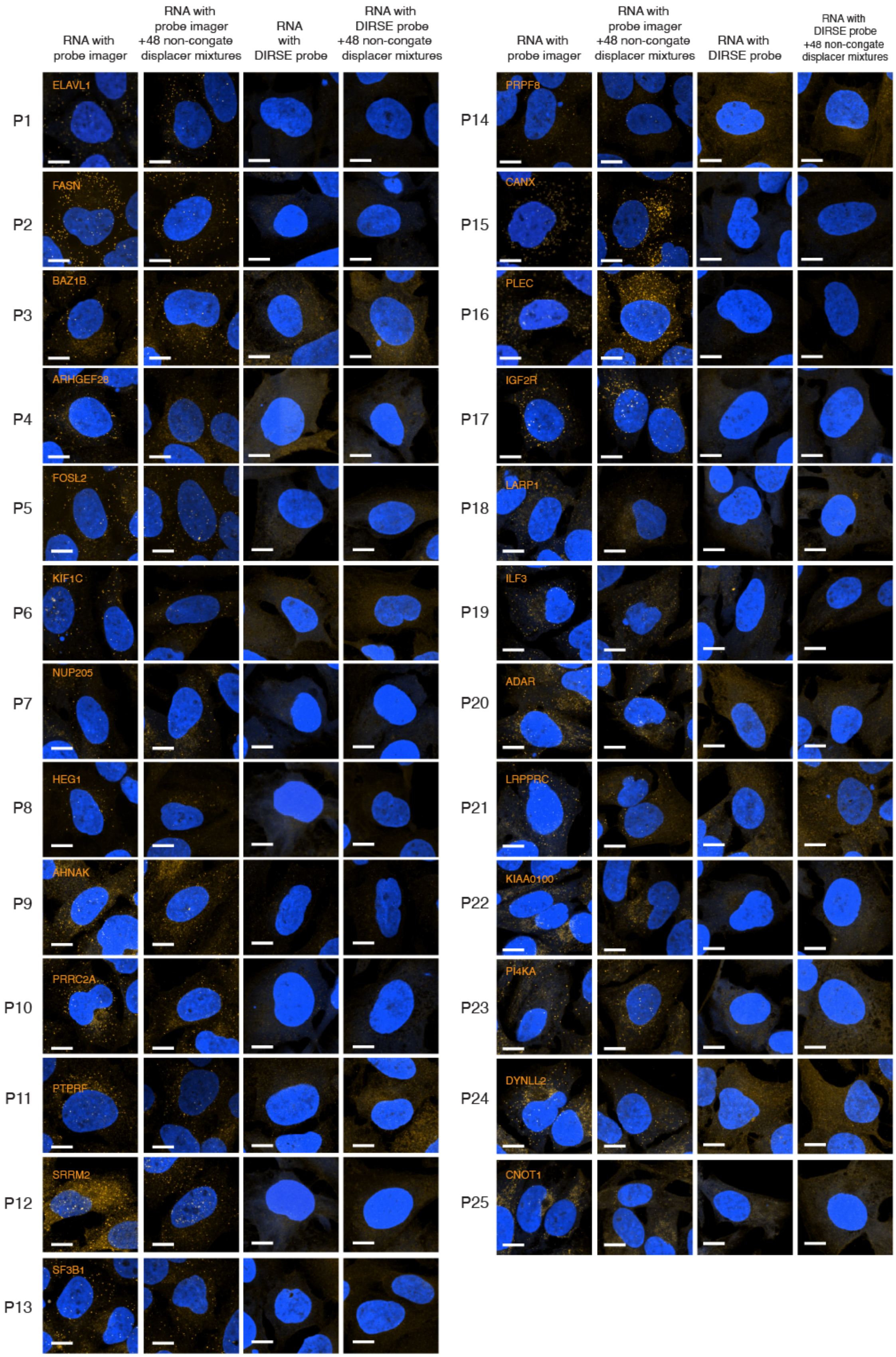

RNA with probe imager
RNA with probe imager +48 non-congate displacer mixtures
RNA with DIRSE probe
RNA with DIRSE probe +48 non-congate displacer mixtures
P1
ELAVL1
P2
FASN
P3
BAZ1B
P4
ARHGEF28
P5
FOSL2
P6
KIF1C
P7
NUP205
P8
HEG1
P9
AHNAK
P10
PRRC2A
P11
PTPRF
P12
SRRM2
P13
SF3B1
P14
PRPF8
P15
CANX
P16
PLEC
P17
IGF2R
P18
LARP1
P19
ILF3
P20
ADAR
P21
LRPPRC
P22
KIAA0100
P23
PI4KA
P24
DYNLL2
P25
CNOT1

**Figure S4. The validation of orthogonality for the designed 25 DIRSE probes.** The fluorescent signal of probe and imager only was measured with the corresponding displacers, and non-cognate 48 other displacers. Non-cognate displacer mixtures didn't induce any significant signal change, indicating high orthogonality between the displacement reactions.

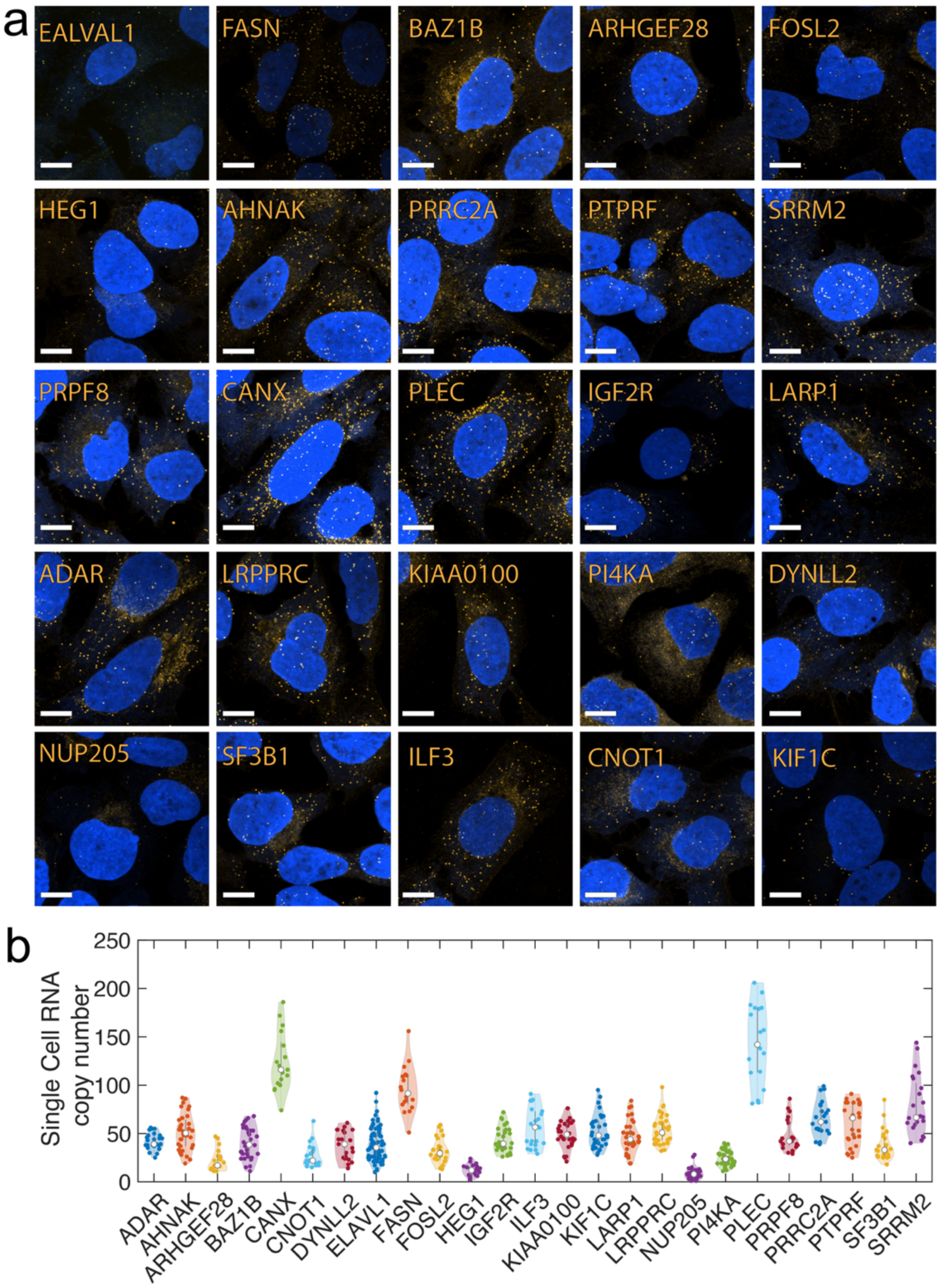

**Figure S5. smFISH imaging for the selected RNA targets in U2OS cells.** (a) The fluorescent imaging of selected 25 RNAs with smFISH. All the scale bars are 10 µm. (b) The violin plots of resolved single cell RNA expression level with smFISH for all the 25 RNA targets. N> 20 cells are used for the statistical analysis.

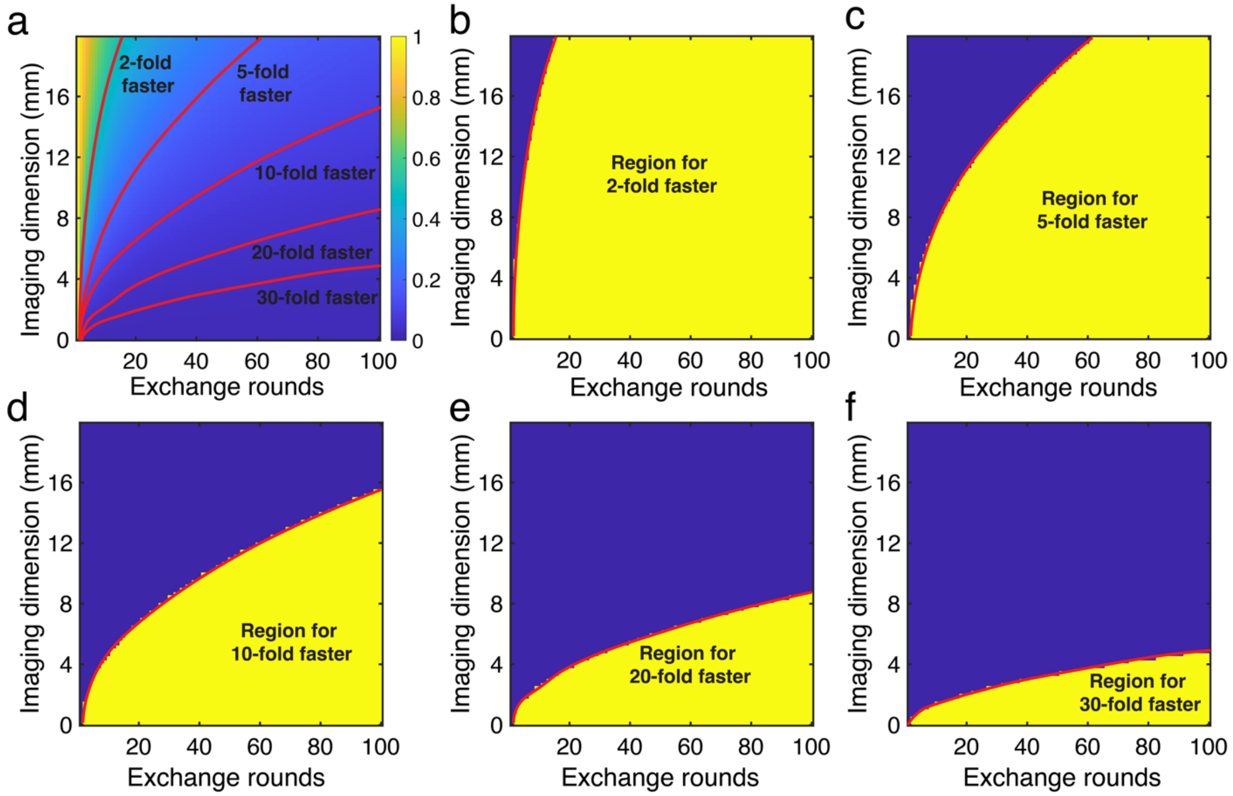


**Figure S6. The comparison of time consumption between the DIRSE and fluidic exchange method.** (a) The heatmap of time consumption ratio of DIRSE to fluidic exchange; (b)(c)(d)(e)(f) The interface indicating the areas that DIRSE workflow is 2-fold, 5-fold, 10-fold, 20-fold, and 30-fold faster than fluidic-exchange workflow, respectively.

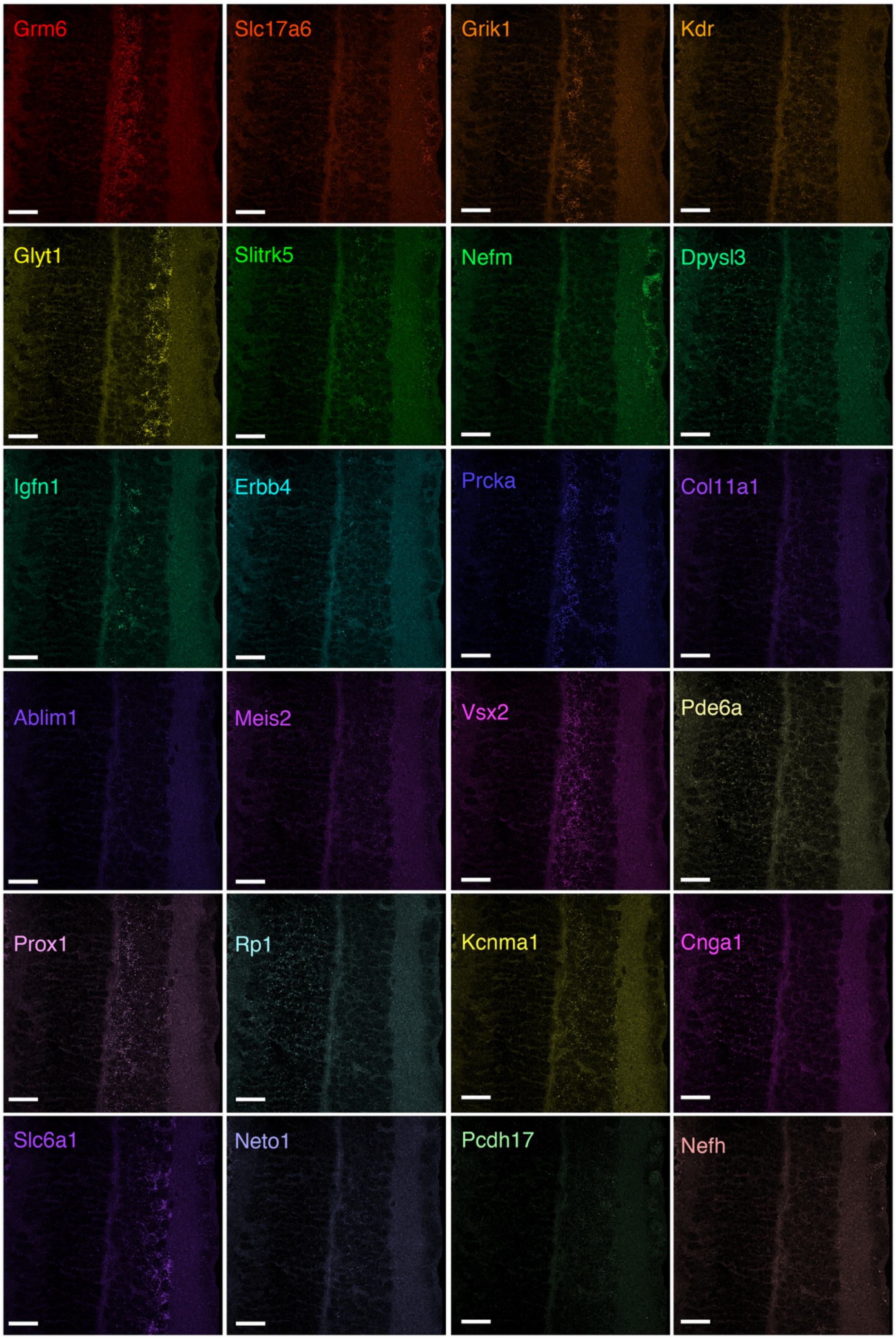


**Figure S7. Fluorescent images of individual 24 RNA target after each round of displacement reactions.** All the 24 RNAs were shown in desired location in the retinal tissues. The scale bars are 20 μm.

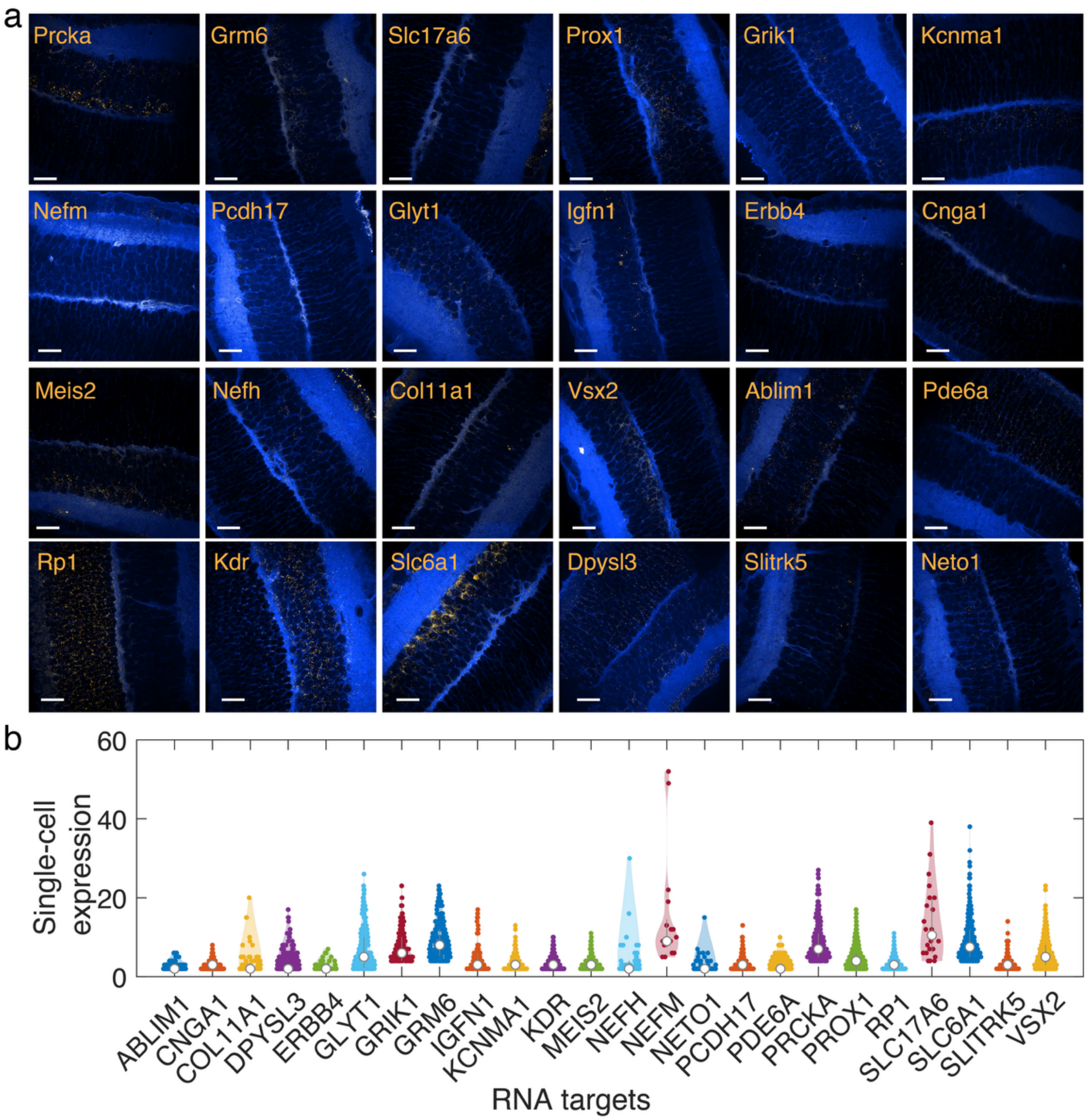


**Figure S8. Single cell expression of RNA with smFISH in in retinal tissues.** (a) Fluorescent images of smFISH for the 24 RNA targets. The cell membrane was stained with WGA. All the scale bars are 20 µm. (b) Violin plots of the RNA expression level in the expressed cell types, n > 20 cells were used for the statistical analysis.

**References.**

1. Stringer, C., Wang, T., Michaelos, M. & Pachitariu, M. Cellpose: a generalist algorithm for cellular segmentation. *Nature methods* **18**, 100-106 (2021).
2. Eng, C.-H.L. et al. Transcriptome-scale super-resolved imaging in tissues by RNA seqFISH+. *Nature* **568**, 235-239 (2019).
3. Chen, K.H., Boettiger, A.N., Moffitt, J.R., Wang, S. & Zhuang, X. Spatially resolved, highly multiplexed RNA profiling in single cells. *Science* **348**, aaa6090 (2015).
